# Feeder cell – the key component in producing scalable and fit NK cells for therapeutic use

**DOI:** 10.64898/2026.04.21.718880

**Authors:** Mira Saari, Farhana Jahan, Lotta Andersson, Anna Syreeni, Pirjo Vehmaan-Kreula, Jan Koski, Emmi Järvelä, Erja Kerkelä, Henrik Paavilainen, Diana Schenkwein, Seppo Ylä-Herttuala, Kim Vettenranta, Helka Göös, Matti Korhonen

## Abstract

Natural killer (NK) cells are increasingly recognized as a versatile therapeutic platform, yet their translation is hindered by limited *ex vivo* proliferation. Feeder cells serve as robust stimulatory component supplying activating signals required to initiate large-scale NK cell expansion. Here, using bench-scale cultures, we evaluated how distinct engineered K562-based feeder cells influence NK cell proliferation, phenotype maintenance, potential for activation, and post-cryopreservation function. Across conditions, feeder-based systems consistently enabled superior, up to 500-fold higher NK cell yield compared to feeder-free system. Variants incorporating membrane-bound costimulatory and cytokine cues yielded the most favorable balance between expansion and functional preservation. Simple adjustments to cryopreservation, including high-density-freezing and centrifuge-free-thawing, further supported NK cell recovery. Together, these findings highlight feeder cells as essential upstream reagents for effective NK cell bioproduction and provide foundational biological insights to guide the rational design and validation of future scalable NK cell manufacturing platforms.

**GRAPHICAL ABSTRACT:** 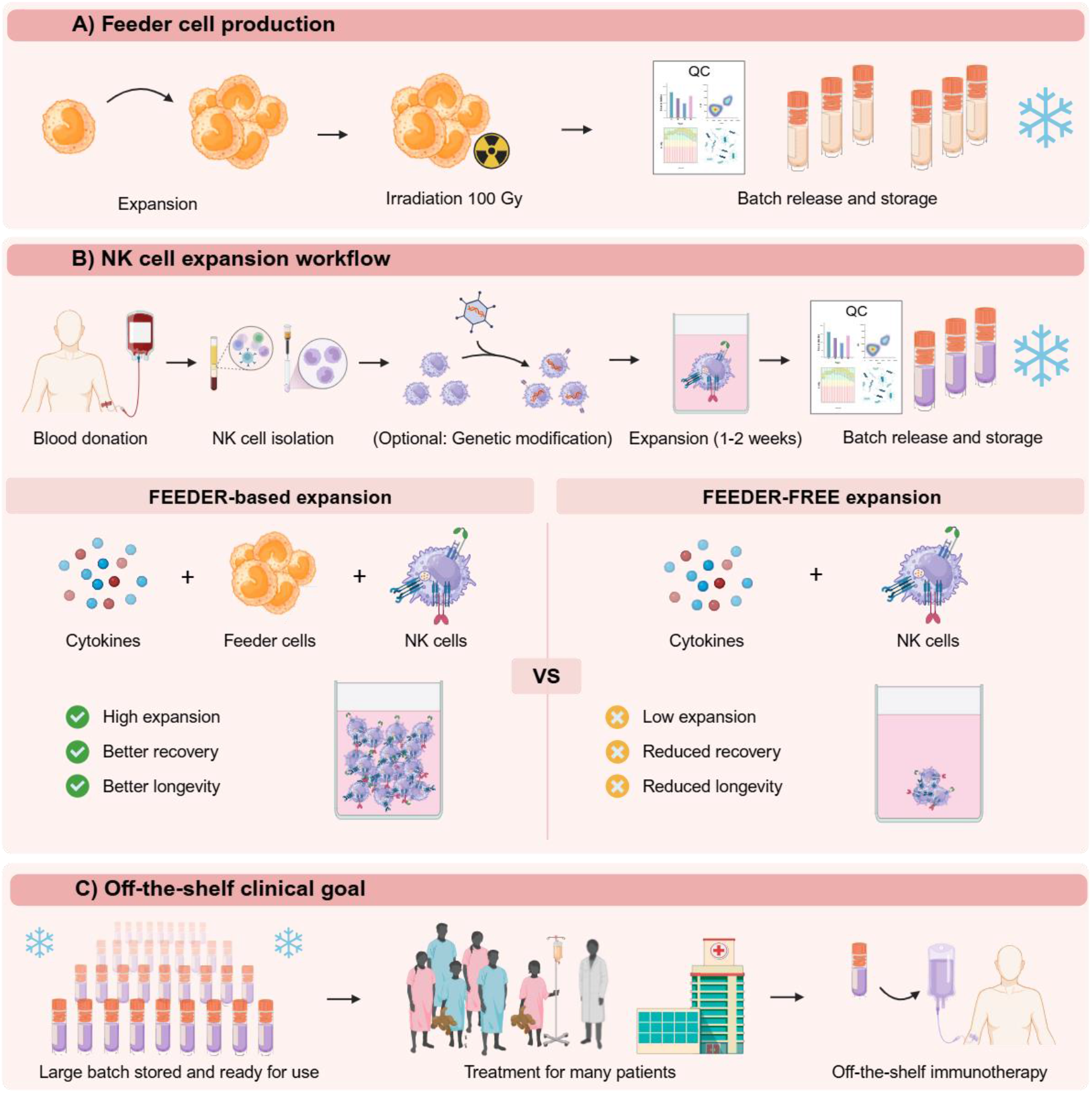

## INTRODUCTION

Natural killer (NK) cells are innate lymphocytes – **effector cells** (see Glossary), with a remarkable ability to recognize and eliminate virally infected and malignant cells without prior sensitization. Their potent **cytotoxicity**, rapid **cytokine** production, and ability to influence **adaptive immunity** make them compelling candidates for cancer **immunotherapy**. Unlike T cells, NK cells carry a low risk of graft-versus-host disease (**GvHD**) and cytokine release syndrome (**CRS**), making them ideal for **allogeneic**, off-the-shelf therapies. However, the broader clinical implementation of NK cells into truly off-the-shelf therapies is limited by slow natural **proliferation** and the modest fold expansion achieved with growth stimulating cytokines alone [1,2].

To boost the therapeutic potential of NK cells, several strategies have emerged. These include engineering NK cells with e.g. chimeric antigen receptors (**CAR**) to improve tumor targeting – often combined with self-induced secretion of supportive cytokines to improve persistence and safety switches to kill any misbehaving NK cells [3,4]. CAR NK cells are showing promising safety and efficacy in early clinical trials for hematologic malignancies, with expanding interest in solid tumors. However, each added manipulation step increases manufacturing complexity, which may reduce overall cell yield due to transduction inefficiency, editing-associated stress, and selection losses [5]. Consequently, maximizing expansion efficiency remains critical for scalable NK cell production. Other less invasive approaches to improve NK cells involve inducing **memory-like** functions using cocktails of **interleukins** such as IL-12, IL-18, IL-15 and IL-21, or sourcing NK cells from **cord blood** to harness their youthful proliferative capacity [6–9]. While all these strategies can boost cytotoxicity and persistence, they fail to overcome the same critical bottleneck: the limited yield of NK cells [10].

Given the relatively low frequency of NK cells collectable in peripheral blood, scalable manufacturing requires efficient *ex vivo* expansion platforms. Among available approaches, **feeder-based** systems consistently achieve high fold amplification, often reaching thousand-to ten-thousand-fold expansion, exceeding typical cytokine-only workflows [1,11]. Although feeder systems are employed in several clinical development programs, their integration into standardized GMP manufacturing workflows remains poor, partly due to step required to gain regulatory approval, and, on the other academic side, the strong focus on improving genetic engineering strategies in current NK cell development [12–15].

However, academic use of feeder cells has long history, particularly with those employing genetically modified **K562** cells (a chronic myelogenous leukemia cell line). They can be engineered to express membrane-bound (**mb**) cytokines such as IL-15 or IL-21, along with **co-stimulatory** ligands like 4-1BBL, CD48, or OX40L, which together promote NK cell proliferation and activation [16–19]. K562 cells are especially well-suited for this role due to their natural expression of multiple NK cell **activating ligands**, including ULBPs, MIC-A/B, CD80, CD86, CD112 and CD155 [20].

The optimal combination of feeder cell modifications – to maximize expansion while preserving NK cell **phenotype, cytotoxicity**, and long-term functionality – remains unresolved. Recent studies have underscored the importance of cytokine and co-stimulatory molecule selection in shaping NK cell fate. IL-15 is widely used to support NK cell expansion, while prolonged exposure may lead to **exhaustion** *[6]*. In contrast, IL-21 has been linked to improved **telomere** maintenance, enhanced safety, and superior anti-tumor activity in engineered NK cells *[7,21]*. A short-term exposure to a combination of IL-12, IL-15, and IL-18 induces a memory-like NK cell phenotype with elevated **IFN-γ** production and long-term persistence [9]. Among activating and co-stimulatory molecules, 4-1BBL:4-1BB engagement is known to drive robust expansion, CD48:2B4 engagement helps prevent NK cell **fratricide**, and properdin:NKp46 interaction enhances activation [22–24].

Based on these findings, we engineered and systematically evaluated a panel of K562 feeder cell variants expressing distinct combinations of 4-1BBL, mbIL-21, mbCD48, mbProperdin, and a membrane-bound cytokine chain of IL-12, IL-18, and IL-21. Then we assessed how feeder-based and feeder-free expansion platforms influence NK cell proliferation, receptor expression profiles, telomere dynamics, and cytokine secretion after NK cells had been expanded and **cryopreserved**. We further assessed how expansion duration influenced long-term proliferation, re-stimulation capacity, and functional competence post-cryopreservation – parameters relevant to manufacturing workflows incorporating cryopreservation storage. Finally, we evaluated cryopreservation strategies by comparing freezing densities, thawing protocols and clinically compatible cryoprotectants to determine their impact on NK cell recovery and viability.

Together, these findings define manufacturing-relevant principles linking feeder cell design, expansion control, harvesting point, and cryopreservation performance in future NK cell production pipelines.

## RESULTS

### Feeder cell engineering minimizes immunosuppressive culture environment

To generate robust K562-based feeder cells, **chimeric transgene constructs** were first designed ***in silico***. We selected 4-1BBL to drive robust NK cell proliferation, mbIL-21 to protect telomeres during rapid proliferation, mbCD48 to prevent NK cell fratricide, mbProperdin to enhance activation via NKp46 signaling, and membrane-bound cytokine chain of IL-12, IL-18 and IL-21 (mbIL12-18-21) to promote memory-like NK cell features (Figure 1A) [7,9,21–24]. K562 feeder cells were **transduced** to express these molecules, delivering tunable stimulatory signals to modulate the NK cell culture environment. Successful surface expression of all constructs was confirmed by **flow cytometry**. The presence of the mbIL12-18-21 construct was detected using antibodies against IL-21 and IL-12p40, where as IL12p35 and IL-18 specific antibodies failed to bind, likely due to **epitope masking** (Figure 1B).

**Figure 1.**
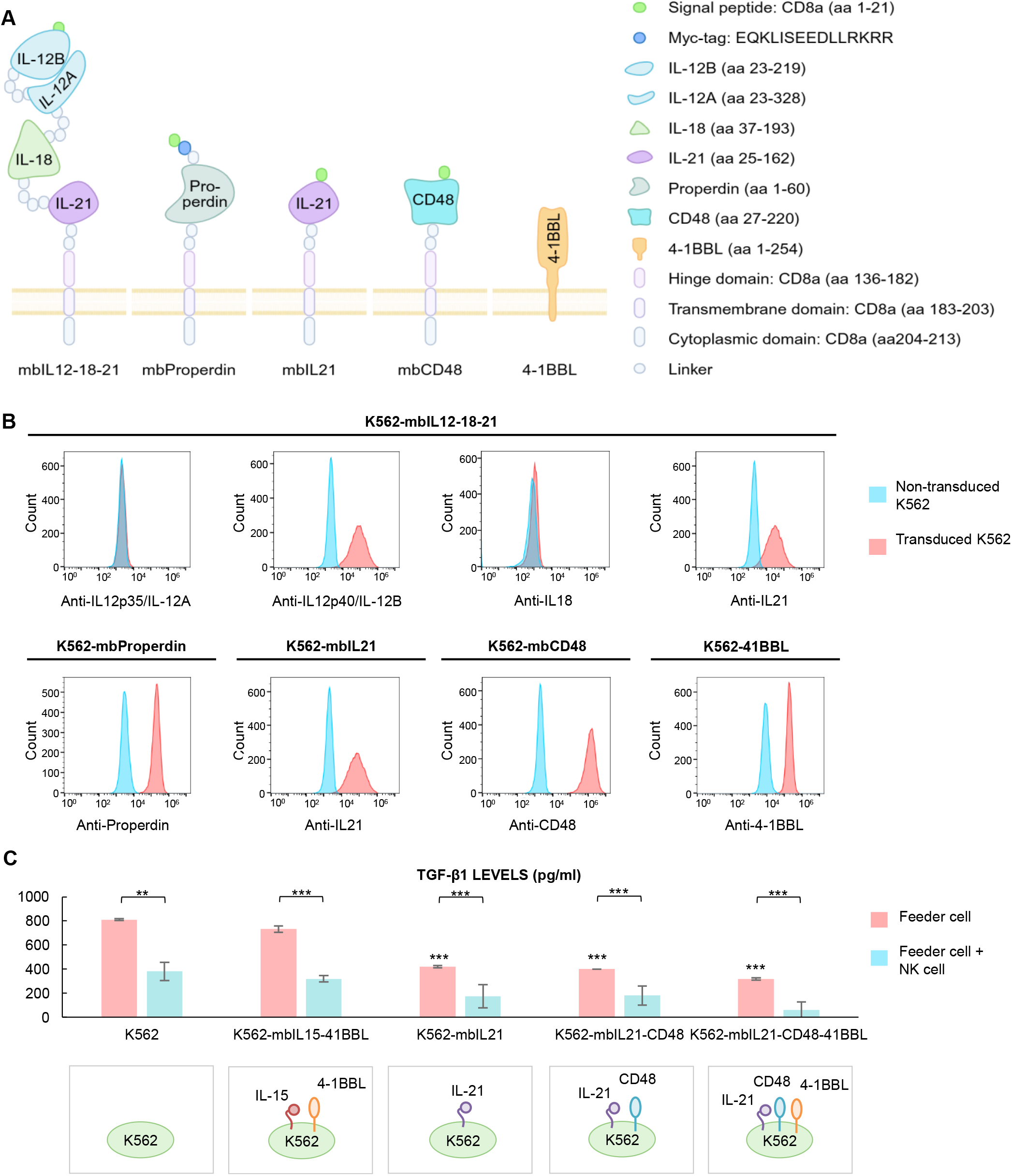
Design and characterization of genetically modified feeder cells to support NK cell expansion. (A) Schematic representation of chimeric transgene constructs introduced into K562 feeder cells via lentiviral transduction. (B) Representative flow cytometry histograms showing surface expression of transgenes following lentiviral transduction of the K562 cell line. (C) Quantification of TGF-β1 from supernatants of irradiated feeder cells (illustrated) cultured alone or co-cultured with NK cells over a 72-hour period. Co-cultures were initiated at a density of 0.5 × 10^6^ cells/mL, with NK cells added at an effector-to-target (E:T) ratio of 1:5. Supernatants were collected at the endpoint. Data are presented as mean ± standard deviation (SD) from n = 3 individual donors. Statistical analysis comparisons of feeder cells were made against K562 cells and another comparison between feeder cells alone vs co-cultures as indicated by lines between the pairs using two-sample paired t-test and two-sample unequal-variance t-test. p < 0.05 (*), p < 0.01 (**), p < 0.001 (***).

Irradiated feeder cells (100 Gy) were cryopreserved as ready-to-use batches to ensure safety and prevent unwanted proliferation. Upon thawing and addition to NK cell expansion cultures, the feeder cells did not proliferate, but maintained their capacity to support NK cell growth, confirming their suitability for batchwise production.

To evaluate the individual and combined effects of the engineered modifications, we generated both singly transduced feeder cell lines (e.g., K562-mbIL21) and combinatorially modified lines (e.g., K562-mbIL21-CD48 and K562-mbIL21-CD48-4-1BBL). Engineered feeder cells exhibited distinct functional profiles. Certain combinations supported high-magnitude NK cell expansion, whereas others produced more selective, lower-magnitude proliferation: a comprehensive list of the generated feeder cell lines and their relative expansion performance is provided in Supplementary Table 1.

To facilitate detailed comparison, all NK cell products were expanded in the same expansion medium (referred hereafter as Medium) and stimulated either without feeder cells, or with **GFP**-tagged and **cell sorted** but otherwise unmodified K562 cells, or with K562-mbIL21, K562-mbIL21-CD48, K562-mbIL21-CD48-41BBL and K562-mbIL15-41BBL cells as low to high-expansion benchmarks.

**TGF-β1** is an **immunosuppressive** cytokine with manifold effects on immune cells, including NK cells [25]. We measured immunosuppressive TGF-β1 cytokine accumulation into culture medium over 72 hours of co-culture with NK cells (Figure 1C). Modified feeders exhibited significantly lower basal TGF-β1 levels compared to unmodified K562 cells, especially when engineered with mbIL21-expressing variants. Among all tested feeder cell variants, K562-mbIL21-CD48-41BBL produced the lowest TGF-β1 levels in NK cell co-cultures.

### 4-1BBL engineered feeder cell support increased NK cell expansion

To ensure a consistent and well-characterized starting population for expansion, NK cells were first isolated from peripheral blood mononuclear cells (**PBMC**) and cryopreserved. Flow cytometry analysis was used to confirm the depletion of non-NK cell populations (Supplementary Figure 1).

We then established a standardized platform to study the effect of the selected feeder cells and feeder-free expansion in Medium on NK cell proliferation and compared expansion timepoints of Day 1, Day 10 and Day 17 (nine experimental conditions total) as illustrated in Figure 2A.

**Figure 2.**
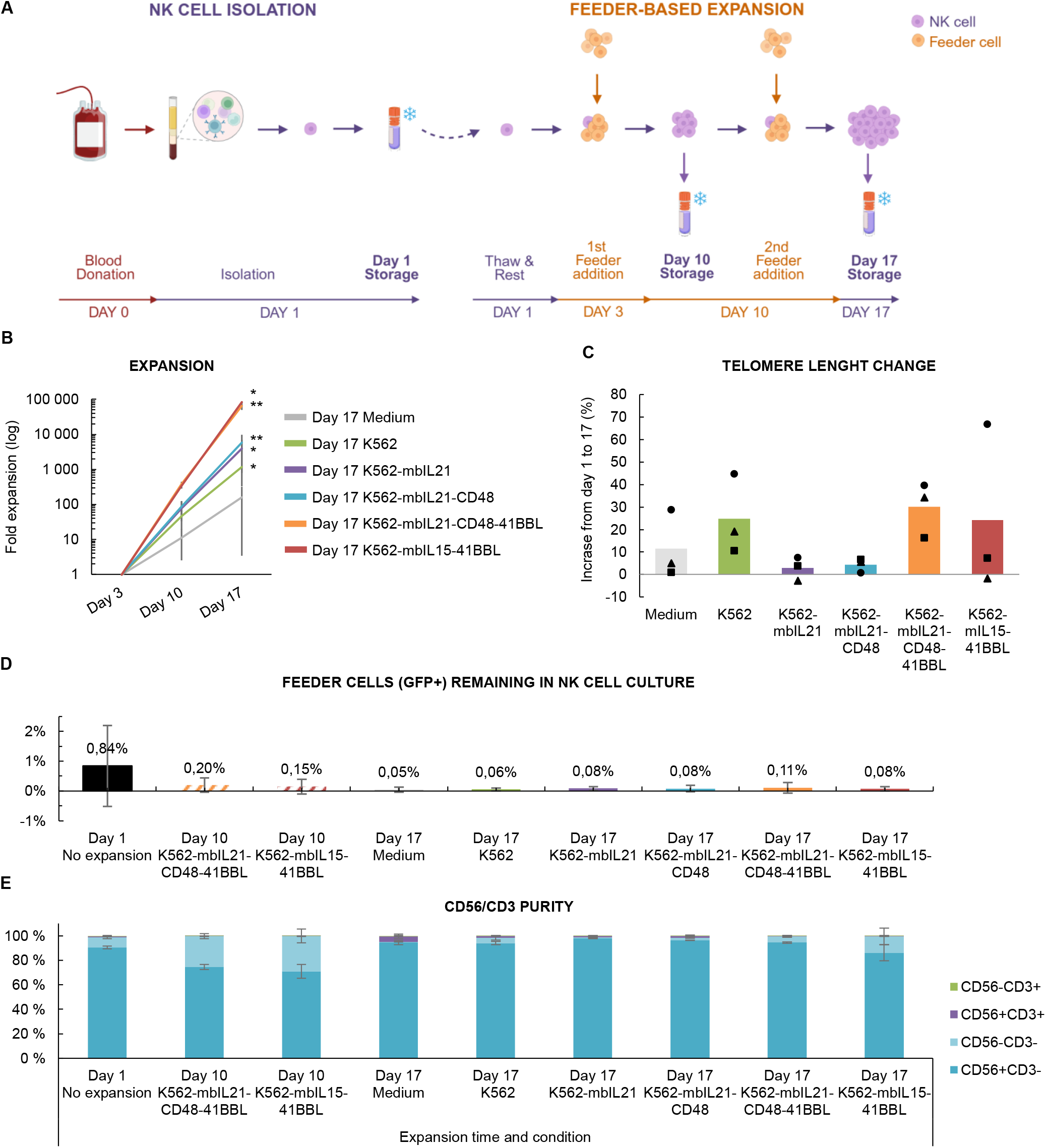
Expansion efficiency and purity of NK cells expanded with or without feeder cells. (A) Schematic overview of the experimental workflow for NK cell expansion, comparing conditions with and without selected feeder cells. The timeline includes the key time points for subculturing and harvesting points. Created with BioRender. (B) Fold expansion of NK cells during expansion with or without feeder cells. (C) NK cells’ telomere length changes from Day 1 to Day 17 using different feeder cells for expansion. DNA was isolated from NK cells at the beginning of the culture (Day 1) and after 17 days of expansion. Telomere length was measured using a PCR-based method, and the relative change over time was calculated for each donor. (D) Flow-cytometry-based assessment of feeder cell clearance (GFP+) from NK cell cultures. (E) Flow cytometry-based assessment of CD56 and CD3 expression to monitor NK cell purity on days 1, 10 and 17 during expansion under various culture conditions, including with and without feeder cells. Data are presented as mean ± SD from (A) n = 6, (B), n = 3 and (C) n = 3 individual donors. Each (C) data point represents an individual donor: donor A (circles), B (triangles) and C (squares). Statistical analysis comparisons were made against the Day 1, non-expanded NK cell condition unless otherwise indicated using two-tailed paired t-test. p < 0.05 (*), p < 0.01 (**), p < 0.001 (***).

Following thawing, NK cells underwent a 3-day recovery phase before feeder addition on Days 3 and 10 at an effector-to-target (**E:T**; NK-to-feeder) ratio of 1:5, with daily subculturing. Because expansion outcomes are highly sensitive to E:T ratios, subculture parameters were optimized for benchtop scale expansion (Supplementary Figure 2). NK cells were harvested for cryopreservation on Day 10 and 17, and all conditions were subjected to downstream analyses including phenotyping, functional assays, telomere length measurement, and long-term culture assessment. Day 10 samples were chosen only from expansions stimulated with the most potent feeder cells.

Among the expansion conditions, the highest NK cell proliferation was achieved by stimulating NK cells twice with either K562-mbIL21-CD48-41BBL or K562-mbIL15-41BBL feeder cells – both expressing 4-1BBL, a ligand associated with enhanced NK cell proliferation (Figure 2B) [26]. NK cells stimulated with K562-mbIL21-CD48-41BBL or K562-mbIL15-41BBL achieved higher total yield and purity compared with Medium cultures, reaching up to 500-fold greater expansion (Supplementary Table 1). K562-mbIL21-CD48-41BBL demonstrated the greatest donor-to-donor consistency. Thus, this feeder cell configuration produced the most significant and reproducible improvements relative to Medium.

To assess the replicative potential of the expanded NK cells, we compared telomere lengths between Day 1 and 17 NK cells. On average, the telomere lengths increase during the expansion 3 – 30% (Figure 2C) without correlation to population doubling times (Supplementary Figure 3). Strongest telomere length gains were observed in NK cells expanded using K562-mbIL21-CD48-41BBL feeder cells. However, telomere dynamics showed considerable donor-specific variability rather than clear feeder-driven trends.

We confirmed the absence of feeder cells in the NK cell cultures by assessing the presence of GFP+ cells. No large GFP+ cells were detected in any of the NK cell cultures. However, small GFP+ events, slightly separated from the main NK cell populations, were observed in all samples, including those derived from cultures that had not been expanded with feeder cells, indicating minor GFP background signal (Figure 2D).

Finally, NK cell purity across all conditions was evaluated using **CD56** and **CD3** expression profiles (Figure 2E). While most conditions maintained high purity, Day 10 samples exhibited reduced CD56 expression compared with Day 1, likely a side-effect of rapid proliferation at this intermediate timepoint as by Day 17, when proliferation slowed, CD56 expression was restored. Notably, although both K562-mbIL21-CD48-41BBL and K562-mbIL15-41BBL feeder cell generated the largest NK cell batches by Day 17, NK cells expanded with K562-mbIL21-CD48-41BBL retained better CD56 expression and overall purity, suggesting a more favorable balance between expansion magnitude and phenotype preservation. A summary of expansion efficiency and population doubling across all tested feeder cell types is provided in Supplementary Table 1.

### Early harvesting improves NK cells’ post-cryopreservation growth

NK cells used in off-the-shelf allogeneic immunotherapies are likely sourced from cryopreserved cell banks. Hence, understanding how expansion strategies influence post-thaw performance is critical for optimizing therapeutic efficacy. To evaluate the post-thaw long-term viability and proliferative potential, expanded and cryopreserved NK cells were thawed and cultured in low cytokine conditions for as long as they remained viable, and weekly cytotoxicity assays were performed to monitor functional persistence.

“The youngest” non-expanded, NK cells survived in post-thaw culture the longest, up to 14-15 weeks (Figure 3A), followed by NK cells expanded with K562-mbIL21-CD48-41BBL. In contrast, expansion with unmodified K562 feeder cells resulted in shortest longevity, suggesting that the addition of cytokines and co-stimulatory ligands to the feeder cells may be necessary to sustain NK cells persistence.

**Figure 3.**
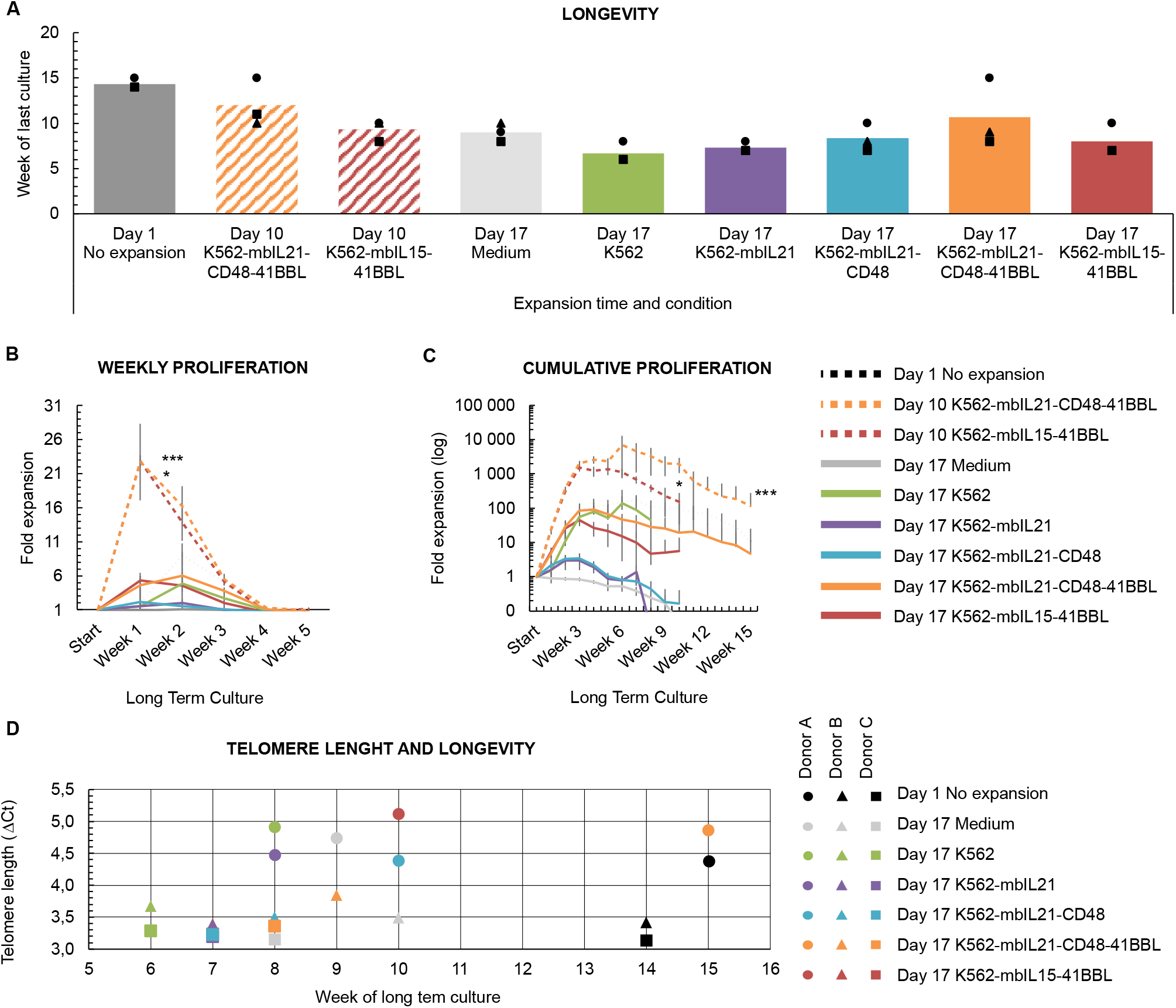
Longevity and proliferative capacity of expanded and cryopreserved NK cells in a long-term ex vivo culture. (A) Post-thaw longevity of NK cells assessed by the number of weeks they remained viable in culture following cryopreservation. The cells were cultured in expansion medium supplemented with 100 IU/ml IL-2. (B) Weekly proliferation and (C) cumulative proliferation curves of thawed NK cells maintained in long-term culture. (D) Relationship between telomere length (ΔCt) and post-thaw longevity of individual Day 1 NK cell samples and those expanded for 17 days. Y axis (ΔCt) shows telomere length expressed as ΔCt values, calculated as the difference between the telomere Ct and the reference gene Ct. Lower ΔCt corresponds to relatively longer telomeres. Data are presented as mean ± SD from (A), (B), (C) and (D) n = 3 individual donors. Data (A) and (D) points represent individual donors: donor A (circles), B (triangles) and C (squares). Statistical analysis comparisons were made against the Day 1, non-expanded NK cell condition unless otherwise indicated using two-tailed paired t-test. p < 0.05 (*), p < 0.01 (**), p < 0.001 (***).

Weekly proliferation analyses revealed that expanded NK cells exhibited highest post-cryopreservation fold expansion during the first week post-cryopreservation, followed by a steady decline until proliferation ceased entirely by week 4 across all conditions (Figure 3B). NK cells expanded with K562-mbIL21-CD48-41BBL or K562-mbIL15-41BBL and harvested on Day 10 displayed most robust proliferation – exceeding 20-fold – whereas NK cells harvested on Day 17 proliferated far less as seen also in the vast variation in NK cell number in **cumulative proliferation curves** (Figure 3C). In comparison, non-expanded NK cells (Day 1) reached their peak proliferation in long-term culture during week 2, consistent with typical behavior of **primary NK cell** cultures.

Importantly, the stimulatory influence of the feeder cells persisted even after cryopreservation. Together these results indicate that NK cells harvested earlier on Day 10 are at their peak proliferative capacity, which could make them ideal for clinical anti-cancer treatment. In contrast, longer culture produces larger total batch sizes, enabling multiple dosing strategy to compensate for loss of post-cryopreservation proliferation.

To test the potential of the long-term-cultured NK cells to respond to a proliferative stimulus, we re-stimulated them on week 8 using K562-mbIL21-CD48-41BBL at E:T of 1:5. Despite having had their proliferation halted since week 4, all studied NK cells demonstrated renewed expansion (Supplementary Figure 4). However, the magnitude of re-expansion (up to ∼1 000-fold) remained substantially lower than the original expansion achieved with first-time-expanded NK cells (>60 000-fold), underscoring the functional limitations of aged NK cells in responding to targets.

Because telomere length may inform on future longevity of therapeutic NK cells, we compared the pre-culture telomere length with post-thaw survival but found no association (Figure 3D). This suggests that donor-specific factors, combined with expansion history, may play a more prominent role than telomere status alone in predicting long-term viability.

### Expansion method influences NK cell cytotoxicity

The capacity of NK cells to eliminate malignant cells is the key feature that makes them valuable in immunotherapies. To assess the long-term functional capacity of expanded NK cells after cryopreservation, we evaluated their cytotoxicity against K562 target cancer cells at multiple timepoints during long-term culture: immediately after thawing (Figure 4A), at week 1 (Figure 4B), week 4 (Figure 4C) and during the week preceding the last survival week of each culture (Figure 4D). NK cells harvested at Day 10 – at strong proliferation stage at the time of cryopreservation – exhibited low cytotoxicity immediately after thawing (Figure 4A). However, one week after thawing, most conditions had regained robust killing activity (Figure 4B). Notably, NK cells expanded with K562-mbIL21-CD48-41BBL or K562-mbIL15-41BBL demonstrated consistently high cytotoxicity through week 4 (Figure 4C). Importantly, even after NK cells had exhausted their proliferation in culture, they retained measurable cytotoxicity in killing assays – demonstrating that effector function can persist even when proliferative capacity is lost, and that the two are not firmly linked.

**Figure 4.**
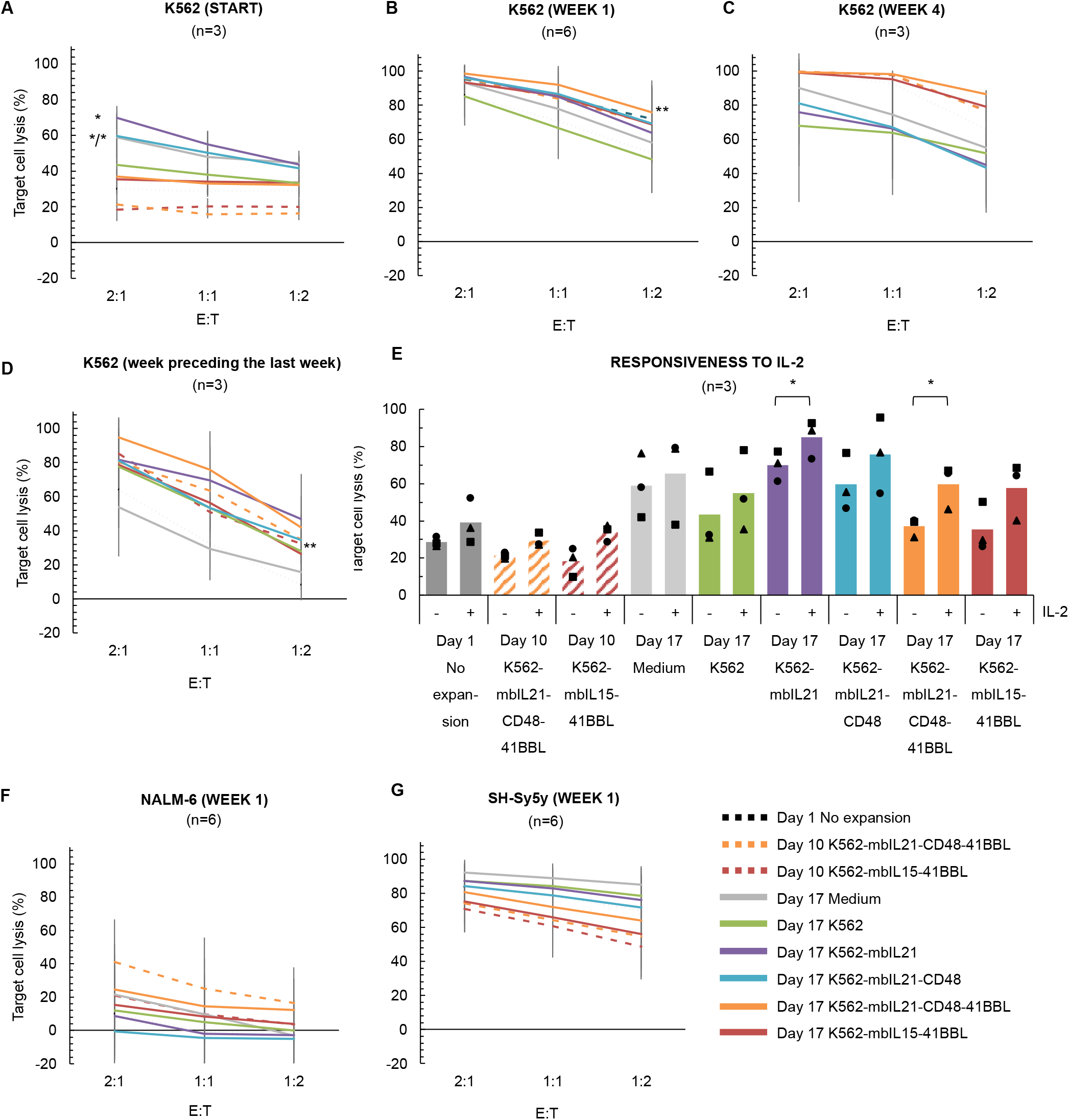
Functional integrity and IL-2 responsiveness of cryopreserved, expanded NK cells. NK cell cytotoxicity against K562 target cells was assessed in a 24-hour co-culture assay at varying effector-to-target (E:T) ratios. NK cells were collected for the assay at multiple timepoints: starting immediately after thawing (A), and at weeks 1 (B) and 4 (C), as well as during each sample’s the week precedin the last week of survival lon -term culture (D). (E) Responsiveness to IL-2 stimulation was evaluated in thawed NK cells using a 24-hour co-culture assay with K562 cells immediately after thawing. 500 IU/ml IL-2 was either added (+) or omitted (–) from the co-culture to assess its effect on cytotoxic activity. (F) Cytotoxicity against NALM-6 cells and (G) SH-Sy5y cells were measured one week after thawing using 24-hour co-culture assays. Each dot represents an individual donor; bars indicate mean values. Data are presented as mean ± SD from (A) n = 3, (B) n = 6, (C) n = 3, (D) n = 3, (E) n = 3, (F) n = 6 and (G) n = 6 individual donors. Data (E) points represent individual donors: donor A (circles), B (triangles) and C (squares). Statistical analysis comparisons were made against the Day 1, non-expanded NK cell condition and in (E) by comparing IL-2 added and omitted condition, using two-tailed paired t-test. p < 0.05 (*), p < 0.01 (**), p < 0.001 (***).

To determine whether freshly thawed NK cells could benefit from a boost of IL-2 prior to cancer cell engagement, we tested their cytotoxicity against K562 targets in the presence or absence of IL-2 (Figure 4E). All NK cell groups responded to IL-2 with enhanced cytotoxicity, supporting the use of IL-2 to rapidly boost functional readiness of therapeutic NK cells prior to patient infusion.

To evaluate activity against additional cancer targets, we examined cytotoxicity toward **NALM-6** (B-cell acute lymphoblastic leukemia cell line) and **SH-Sy5y** (neuroblastoma cell line) (Figure 4F and G). Even at the peak of post-thaw cytotoxicity, NK cells showed lower killing of NALM-6 than K562, likely reflecting resistance mechanisms characteristic of this cell line. NK cells expanded with K562-mbIL21-CD48-41BBL exhibited the strongest responses among all conditions.

In contrast, NK cells expanded using feeder cells providing more moderate proliferative stimulation showed superior killing of SH-Sy5y cells, suggesting that moderate stimulation can yield functional NK cell products for use against certain cancers. Together, these results show that feeder cell design influences not only expansion magnitude but also the quality and specialization of cytotoxic responses, further supporting the need to tailor feeder cell engineering to the particular therapeutic indication. These differences in killing capacity may be linked to activation or exhaustion profiles caused by different feeder cells, which we further explored below in phenotyping, **degranulation**, and **cytokine secretion** analyses [27].

### Minimally expanded NK cells have favorable surface markers and cytokine profiles

To investigate the activation status and effector readiness of the expanded NK cells, we analyzed changes in **surface expression** of **CD69** (activation marker), **CD107a** (cytotoxic degranulation marker), and the apoptosis-inducing ligands **FasL** and **TRAIL** following target cell encounter (Figure 5A, Supplementary Figure 5A). CD69 expression was elevated across all conditions, likely reflecting prior cytokine exposure during post-thaw culturing *[28]*. However, minimally expanded NK cells exhibited slightly higher CD69 expression, particularly upon target cell encounter. While the frequency of CD107a expression was comparable across groups, the median fluorescence intensity (**MFI**) was higher in minimally expanded NK cells – such as those cultured in Medium alone – suggesting more pronounced degranulation by individual cells. Interestingly, Day 10 NK cells exhibited similar degranulation upon target cell encounter, but higher baseline degranulation, compared to other timepoints (Supplementary Figure 5B). FasL and TRAIL expressions followed a similar trend, with modest upregulation upon target cell stimulation and slightly higher expression in minimally expanded NK cells.

**Figure 5.**
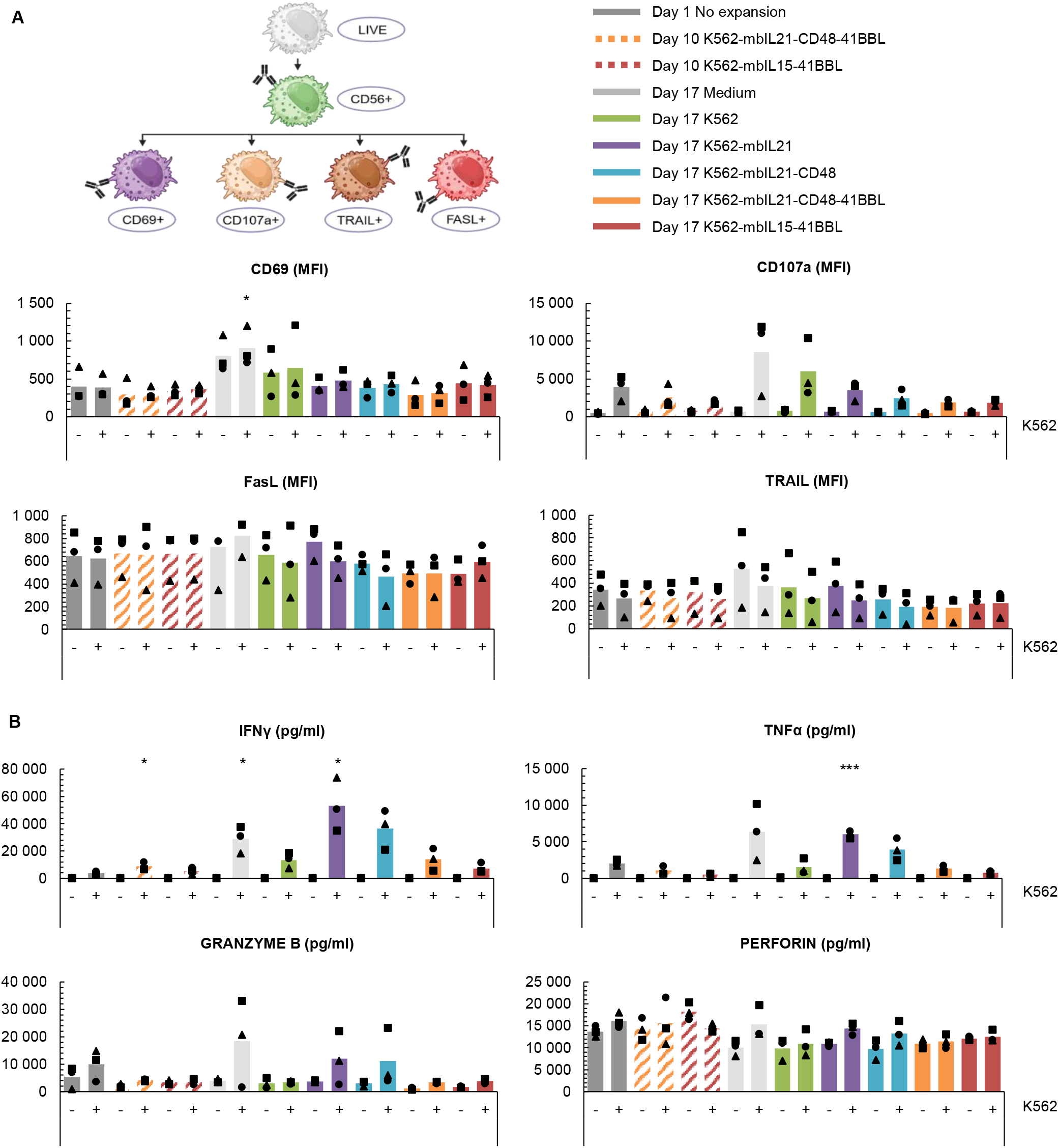
Activation marker expression and cytotoxic effector molecule secretion of NK cells in response to K562 target cells. (A) NK cells were co-cultured for 4 hours with (+) or without (–) K562 target cells and analyzed by flow cytometry after live/CD56+ gating. Surface expression of the activation marker CD69, degranulation marker CD107a, and apoptosis-inducing ligands FasL and TRAIL were measured on NK cells. Marker expression was quantified using median fluorescence intensity (MFI). (B) Secretion of inflammatory cytokines (IFN-γ and N -α) and cytotoxic granule components (granzyme B and perforin) were measured from co-culture supernatants using a multiplex bead-based assay. Data are presented as mean ± SD from (A) and (B) n = 3 individual donors. Data (A) and (B) points represent individual donors: donor A (circles), B (triangles) and C (squares). Statistical analysis comparisons were made against the Day 1, non-expanded NK cell condition unless otherwise indicated using two-tailed paired t-test. p < 0.05 (*), p < 0.01 (**), p < 0.001 (***).

To assess the functional responsiveness of cryopreserved NK cells, cytokine and **cytotoxic granule** secretion were measured using a **multiplex bead-based assay** (Figure 5B). Upon target cell encounter, the secretion of IFN-γ and **TNF-α** increased across most groups, with highest levels detected again in NK cells expanded minimally with Medium or K562-mbIL21 feeder cells. **Granzyme B** secretion was also elevated in these groups, indicating a strong effector profile. **Perforin** levels were high even without stimulation and increased only modestly upon target encounter, consistent with previous reports showing that feeder cell-expanded NK cells maintain an elevated baseline perforin *[29]*.

Collectively, these results show that lower proliferation during NK cell expansion step yields NK cells with superior effector signatures – including enhanced granzyme/perforin readiness and stronger IFN-γ and N -α secretion – despite their reduced expansion rates. Although these activation and effector markers are commonly used to predict NK cell cytotoxicity, their expression did not consistently align with observed killing performance. Here, marker expression patterns suggest Medium, unmodified K562 or K562-mbIL21 cells as top candidates for producing cytotoxic NK cells, which correlated with their immediate post-thaw cytotoxicity against K562, (Figure 4A), and their performance against SH-Sy5y (Figure 4G). However, these markers did not explain differences in long-term cytotoxicity against K562 or NALM-6, where the NK cells expanded with these feeder cells exhibited weakest responses (Figure 4 B-D and F). These marker-to-function disconnects suggest that moderately stimulated NK cell products – those expanded with lower-intensity feeder cells – may align better with certain therapeutic indications where rapid, high-quality cytotoxic responses are essential.

### Feeder cells modify NK cells’ phenotypic profile

As activation markers did not provide reliable predictive information on the true functional ability of the expanded NK cells, we next examined if phenotype markers could be used to evaluate the therapeutic potential of the produced NK cells. We analyzed live cells for surface markers associated with activation, cytotoxicity, and **senescence** one week after thawing (Figure 6A). Flow cytometry analysis showed that CD56 expression was markedly reduced in NK cells harvested on Day 10, aligning with earlier purity data (Figure 2E).

**Figure 6.**
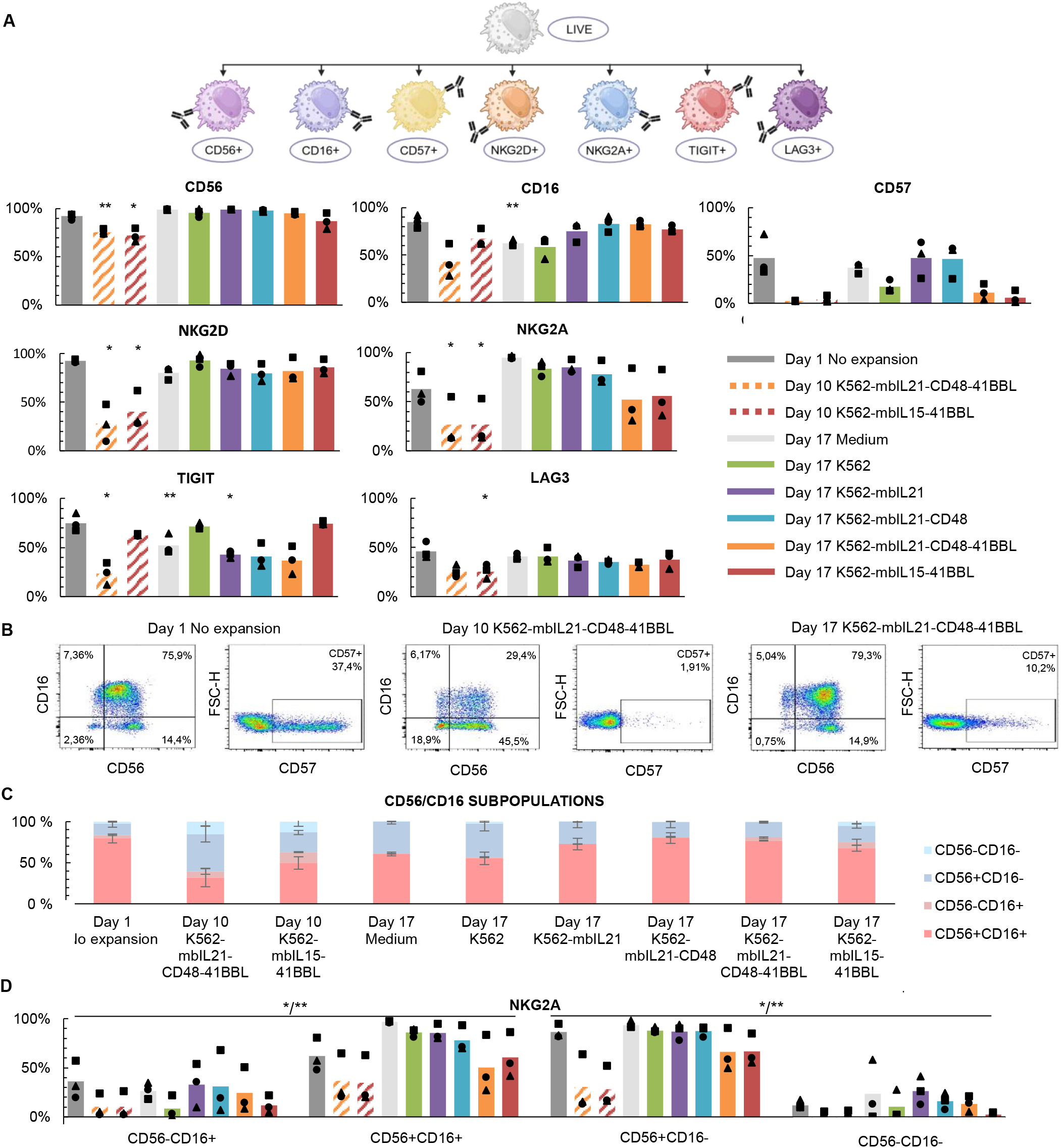
Phenotype changes in NK cell population during expansion with feeder cells. (A) Flow cytometry analysis of NK cell surface markers associated with activation, cytotoxicity, and senescence present in the whole live cell culture population. Expression levels of CD56, CD16, CD57, NKG2D, NKG2A, TIGIT, and LAG-3 were measured from live gating. (B) Representative flow cytometry plots showing changes in CD56/CD16-defined NK cell subpopulations and CD57 expression in NK cells that were either unexposed to feeder cells or expanded with feeder cells. (C) Chart summarizing the relative proportions of CD56/CD16-defined NK cell subsets across all conditions, based on flow cytometry analysis. (D) Chart showing changes in NKG2A expression on CD56/CD16-defined NK cell subpopulations in NK cells that were either unexposed to feeder cells or expanded with feeder cells. Data are presented as mean ± SD from (A) n = 3, (C) n = 3 and (D) n = 3 individual donors. Data (A) and (E) points represent individual donors: donor A (circles), B (triangles) and C (squares). Statistical analysis comparisons (A) were made against the Day 1, non-expanded NK cell condition and (D) were made between CD56^−^CD16^+^vs. CD56^+^CD16^+^, and CD56^+^CD16^−^ vs. CD56^−^CD16^−^ populations, using a two-tailed paired t-test. p < 0.05 (*), p < 0.01 (**), p < 0.001 (***).

**CD16**, a key receptor for antibody-dependent cellular cytotoxicity (**ADCC**) related applications, was highly expressed in Day 1 NK cells, but notably reduced in Day 10 group and in cells expanded with only Medium or unmodified K562 feeder cells. In contrast, NK cells expanded with any of the engineered feeder lines retained CD16 expression, underscoring how feeder engineering can maintain ADCC-relevant features in the final product.

**CD57**, a marker of terminal differentiation and improved cytotoxic potential, was upregulated in NK cells expanded with mbIL21-expressing feeder cells and downregulated by 4-1BBL-expressing feeder cells. The elevated expression of CD57, strong secretion of IFN-γ, TNF-α and granzyme B were all characteristics of minimally proliferating NK cells with high immediate cytotoxicity as seen in K562 cytotoxicity data (Figure 4A). Conversely, CD57 downregulation seen in NK cells expanded with 4-1BBL expressing feeder cells shifted NK cells toward a more proliferative and potentially longer-lived phenotype – consistent with long-term culture data (Figure 3A).

**NKG2D**, a key activating receptor for tumor cell recognition, was highly expressed under most conditions except on NK cells harvested on Day 10 – reflecting decrease in cytotoxicity during the high-proliferation phase. Among **inhibitory receptors**, which can prevent NK cells from performing their cytotoxic functions, **NKG2A** expression was reduced in NK cells expanded with 4-1BBL-expressing feeder cells, while remaining elevated in other conditions. Expression of inhibitory **TIGIT** was lowest in NK cells expanded with mbIL21-expressing feeder cells and highest in those expanded with K562-mbIL15-41BBL. **LAG-3** expression remained low across all conditions but showed a notable decrease in NK cells harvested on Day 10, suggesting a transient regulatory shift during early expansion.

Further analysis into CD57 and CD56/CD16-defined NK cell subpopulations revealed dynamic shifts in population compositions across the expansion timeline, likely contributing to the transient dip in cytotoxicity around Day 10 (Figure 6B, 6C and Figure 4A). Examination of CD56/CD16 subpopulations also revealed connection between CD56+ and NKG2A+ populations, suggesting a link between CD56 expression and NKG2A regulation. (Figure 6D). This observation aligns with previous findings that CD56^bright^ NK cells often co-express NKG2A during early activation, pointing to association between the proliferative or less differentiated NK cell states [30]. Interestingly, these shifts did not uniformly predict cytotoxicity across multiple targets (Figure 4A-G).

Together, these findings illustrate how feeder cell engineering can be used to tune the expression of activating receptors (e.g., NKG2D), suppress inhibitory receptors (e.g., NKG2A, TIGIT), and maintain the ADCC-critical CD16.

### Optimized cryopreservation and thawing protocol improve NK cell recovery

NK cells are sensitive to freezing and thawing, and the recovery of therapeutic NK cells after cryopreservation impacts their proliferation and functionality. Hence, their off-the-shelf therapeutic potential may be impacted by the conditions used during freezing and thawing [31]. To identify optimal parameters for recovery, we cryopreserved CAR NK cells that had been expanded with K562-mbIL21-CD48-41BBL feeder cells, at two densities using different ***in vitro-*** and ***in vivo***-compatible cryopreservation solutions and thawing strategies.

Consistent with our expectations, the number of cells recovered was lower than the number frozen across all methods tested and centrifugation further reduced cell numbers (Figure 7). However, differences between conditions were modest immediately after thawing, with more pronounced differences emerging three days post-thaw. Across all cryopreservation solutions, the recovery of CAR NK cells cryopreserved at higher density was superior compared to those frozen at lower density, particularly when cells were not washed and centrifuged during thawing process. CAR NK cells frozen at high density and thawed without centrifugation not only recovered but proliferated beyond initial counts under both *in vivo-* and *in vitro* -compatible options. Notably, MACS Freezing solution at 50×10^6^ cells/mL yielded the hi hest recovery and most efficient proliferation. Interestingly, a 5% DMSO concentration in Albunorm resulted in higher recovery than 10% DMSO and provided the greatest recovery across all in vivo–compatible formulations. These improvements represent simple, scalable handling adjustments that are easily incorporated into GMP workflows, offering immediate translational benefit for commercial NK cell manufacturing.

**Figure 7.**
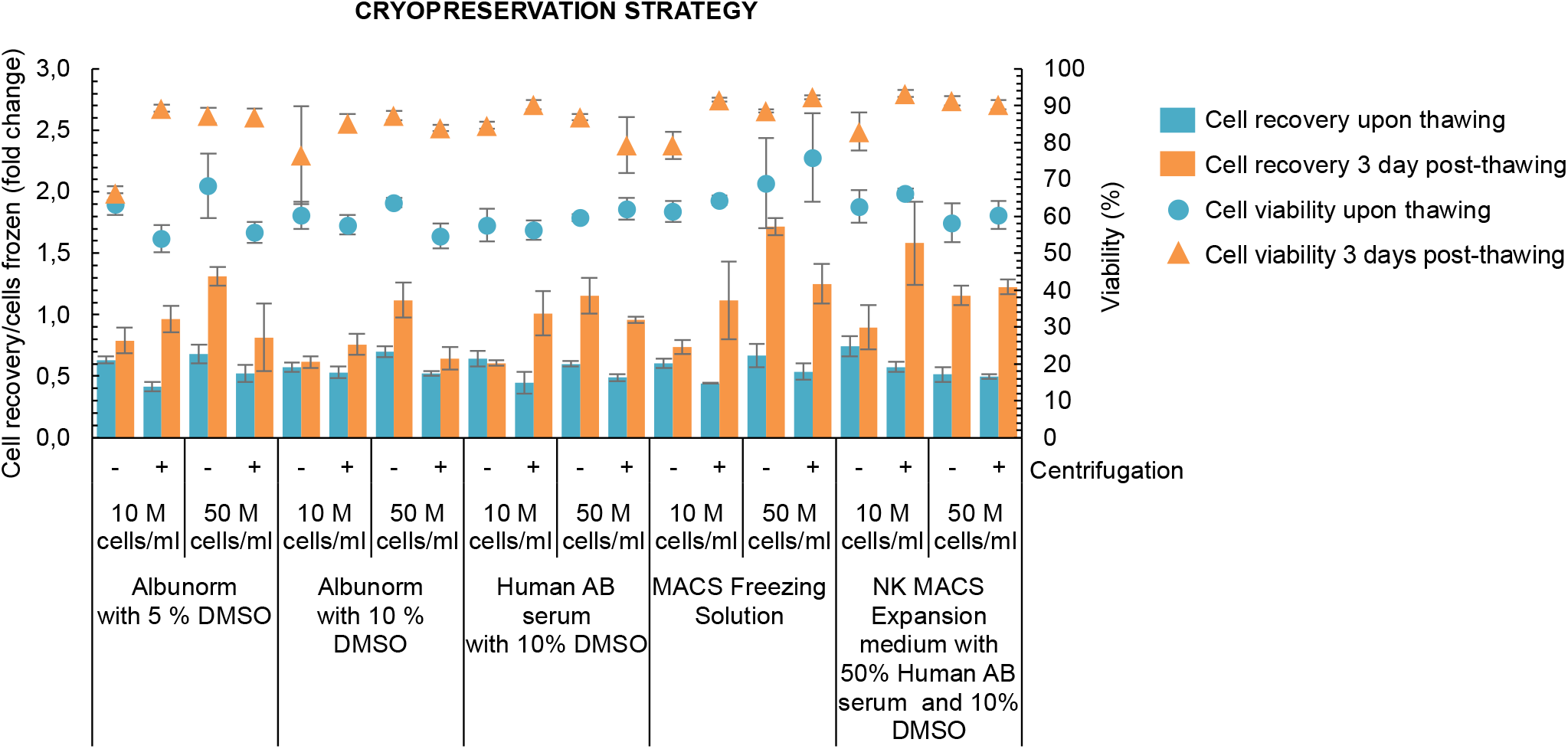
Comparison of cryopreservation methods for expanded NK cells. The NK cells were cryopreserved using five different cryopreservation media at two cell densities, and either with (+) or without (–) centrifugation following thawing. Cells were cultured in NK MACS Expansion Medium, and both cell recovery (expressed as fold change) and viability were assessed immediately upon thawing and three days post-thaw. Fold change in cell number is shown as bars, and viability as overlaid dots. Fold change was calculated relative to the number of NK cells originally frozen per vial, providing an estimate of cell loss during the freeze–thaw cycle and potential recovery during short-term culture. Data are presented as mean ± SD from 3 experiments.

## DISCUSSION

### Feeder-based expansion improves scalability

Efficient large-scale expansion of NK cells remains a major challenge in the development of off-the-shelf immunotherapies. Unlike autologous T cell products, NK cell therapies rely on banking of products made from cell donated by healthy donors. Unfortunately, the naturally low abundance of NK cells collectable from peripheral or cord blood combined with low viability after genetic engineering to produce e.g. CAR NK cells, makes maximizing yield during production essential [5,10,32]. Achieving exponential product amplification is something feeder-based systems offer a distinct bioprocessing advantage for, while also preserving key quality attributes and functional integrity of NK cells [33].

In this study, we systematically compared feeder-free and feeder-based expansion approaches. By evaluating five K562-derived feeder cell variants, we identified feeder cell configurations that most effectively supported robust NK cell proliferation while preserving functional and phenotypic integrity after cryopreservation. Among these, the feeder cell line expressing mbIL21, CD48, and 4-1BBL consistently yielded the strongest overall performance, making it a candidate for master cell banking. However, each feeder variant offered distinct benefits, underscoring the challenge of developing a single “universal” feeder cell suitable for all therapeutic contexts.

This study aimed to systematically assess how distinct engineered feeder cell variants shape NK cell expansion dynamics, phenotypic stability, and post-thaw performance in the context of scalable manufacturing. We decided to use the K562 cell line – one of the most widely used platforms despite their known secretion of the immunosuppressive cytokine TGF-β1 – as a backbone [34]. TGF-β1 inhibits ADCC and suppresses CD16-mediated IFN-γ production in NK cells, making it an undesirable feature in large-scale expansion systems, which is why we also needed to assess if TGF-β1 secretion poses risks [35,36]. Among the feeder lines we tested, the mbIL21-expressing variant secreted less TGF-β1, and NK cells expanded with K562-mbIL21-CD48-41BBL not only proliferated more robustly but also required more frequent medium replenishment, which further dilutes TGF-β1 over time. Despite transient exposure to this inhibitory cytokine, feeder-expanded NK cells retained stronger functional competence than NK cells expanded without feeders. Together, these results highlight the value of comprehensive feeder cell characterization and show that their composition can be purposefully engineered to create a supportive bioprocess milieu that promotes proliferation and minimizes suppressive signals.

Because the expansion phase is when feeder cells contribute most significantly to NK cell manufacturing, we first examined how different feeder configurations influence proliferative capacity. Our results underscored the central role of 4-1BBL -mediated stimulation in achieving high-magnitude NK cell proliferation. Using K562-mbIL21-CD48-1BBL and K562-mbIL15-41 L, we o served 5 000-to 000-fold NK cell expansion within two weeks – levels consistent with previous reports showin 00 000-to 500 000-fold expansion withing three to four weeks [26,37]. Lower 900-fold expansion trends have been reported for cord blood NK and CAR NK cells expanded with K562-mbIL21-CD48-1BBL feeder cells [38]. Also, the inclusion of CD48 into feeder cell design may have further boosted proliferation. Through 2B4 engagement, CD48 can enhance NK cell activation, reduce fratricide and apoptosis, and support more sustained growth while enhancing anti-tumor activity and infiltration into solid tumors [23,38,39].

The expansion rates achieved with our bench top protocol surpass those reported in many clinical trials, where fold expansions typically range from 1 000 to 2 000 over 14–21 days [40]. Theoretically, the protocol developed in this paper using K562-mbIL21-CD48-41BBL expressing feeder cells could expand NK cells to dose numbers 500 times higher than in Medium alone. In this context, feeder-enabled NK cell amplification substantially improves changes of meeting dose requirements for off-the-shelf NK products, depending on the respective clinical indication and dosing strategy [41].

Use of feeder cells for expansion also led to final products with lesser contamination by T and NKT-like cells, possibly by selective stimulation of NK cells over the other populations. We also noted a transient downregulation or shift in intensity of CD56 expression, particularly at intermediate time points during rapid proliferation. This phenomenon, also reported by others, suggests that pairing CD56 with additional markers such as activating receptors may provide a more reliable and standardized quality readout during early harvesting [26].

These combined observations indicate that specific feeder cell engineering as a practical bioprocess lever that influences both expansion magnitude and culture purity. They also underscore the need for bioreactor systems capable of supporting rapid NK cell proliferation under scalable manufacturing conditions.

### Feeder-based expansion enhances characteristics amenable to therapeutic applications

Beyond promoting proliferation, feeder-based expansion shaped NK cell phenotypes relevant to therapeutic application. Although NK cell surface receptor expression alone cannot fully predict cytotoxic performance [42], feeder-expanded NK cells consistently expressed high levels of key activating receptors, including NKG2D and CD16. These features are particularly relevant from a biotechnology perspective, as they support combination strategies with monoclonal antibodies and other targeted cancer treatments [43–46]. NK cells expanded in feeder-free Medium had higher frequencies of CD57, suggesting a trade-off between feeder-based expansion magnitude and feeder-free maturity [47].

Analysis of inhibitory receptor expression further demonstrated that feeder-based expansion can influence NK cells’ therapeutic suitability. In particular, NK cells cultured with K562-mbIL21-CD48-41BBL and K562-mbIL15-41BBL showed substantially lower expression of NKG2A than feeder-free expanded cells. This decrease is most notable in NK cells harvested on Day 10, consistent with emerging evidence of alternative NK cell differentiation pathways [48]. Similarly, expression of inhibitory TIGIT and LAG-3 was reduced on NK cells expanded with all mbIL21-engineered feeder variants. Reversely, high TIGIT was observed in NK cells expanded with K562-mbIL15-41BBL feeder cells, which may be a sign of exhaustion due to overstimulation with mbIL15 and soluble IL-15 supplement [6,49,50].

Collectively, these findings highlight that feeder cell engineering can influence NK cell receptor expression profiles associated with responsiveness to inhibitory signaling within the tumor microenvironment. Although direct resistance was not assessed, modulation of inhibitory receptors such as NKG2A may have implications for therapeutic positioning. For example, high NKG2A expression has been linked to reduced activity against HLA-E^+^tumors but may also identify NK cell products that could benefit from combination with NKG2A-blocking checkpoint inhibitors to restore function [30,51].

### Functional assessment of NK cells requires target-specific evaluation

Cytotoxicity is the defining, functional attribute of therapeutic NK cells and widely used as a primary indicator of their *in vitro* potency. To assess how expansion strategy influences functional performance, we evaluated NK cell cytotoxicity at multiple time points and compared these data with activation related receptor expression, degranulation, and cytokine-secretion profiles. This approach allowed us to identify functional correlations relevant to NK cell product characterization.

NK cells’ immediate post-thaw cytotoxicity against K562 leukemia targets, and later against SH-Sy5y neuroblastoma targets, was strongest in NK cells characterized by elevated CD57 and CD107 expression, robust secretion of IFN-γ, N -α, and ran yme. These NK cells originated from less intensive expansion conditions lacking 4-1BBL stimulation, suggesting that slower expansion kinetics may preserve certain mature effector traits critical in NK cells recovering from cryopreservation or in targeting specific cancer cells. NK cells rapidly expanded with K562-mbIL21-CD48-41BBL or K562-mbIL15-41BBL feeders demonstrated lower immediate cytotoxicity against K562 regardless of high expression of activating cytotoxic receptors. This may reflect NK cells’ need for a transient recovery phase post-cryopreservation or be a sign of temporary functional exhaustion. It may also be a sign of NK cells’ focus on proliferation instead of cytotoxicity, as their post-thaw proliferation speed was several folds higher than that of the other cell preparations. Whichever the reason for low immediate cytotoxicity, one week later, NK cells expanded with K562-mbIL21-CD48-41BBL exhibited a strong cytotoxicity against K562 and NALM-6 leukemia targets.

These findings indicate that no single functional marker reliably predicts cytotoxicity across all tumor types, and that functional responses are inherently recovery and target-dependent [52]. While *in vitro* functional assays remain central to NK cell product characterization, their predictive value may depend on assay timing and tumor target selection. Our studies highlight two manufacturing-relevant considerations. Firstly, NK cell product characterization should include both early and delayed cytotoxicity assessments to capture post-thaw recovery dynamics and avoid under- or over-estimating potency [51]. Secondly, target-specific readouts are required, as NK cell functional hierarchies may vary between cancer targets. Rational refinement of feeder cell design may provide a strategy to align NK cell products with defined therapeutic applications.

### NK cells’ harvesting timepoint defines final batch size

From a manufacturing perspective, harvest timing of NK cells represents an important bioprocess decision point. Earlier harvests yield smaller lots but preserve higher proliferative potential, whereas later harvests lower cost per dose through larger batch sizes but may require a higher number of administered doses due to reduced proliferative reserves. Thus, while achieving high NK cell yields is essential for clinical scalability, preserving the longevity of the final product is equally critical for therapeutic performance. To examine this trade-off, we assessed telomere length, post-cryopreservation proliferation, and long-term responsiveness.

Prolonged *ex vivo* expansion can drive exhaustion and senescence. This is typically reflected in telomere shortening and diminished responsiveness [6,53]. However, we observed no direct association between telomere length and NK cell longevity in our system. Although we hypothesized that mbIL21-engineered feeder cells would promote telomere elongation [7,38], NK cells expanded with all feeder cells showed telomere preservation or telomere length increase. Perhaps feeder cell stimulation is able to activate telomerase (TERT) pathways, supporting continued proliferation without inducing senescence [26,54].

Long-term culture provided additional insight into how harvest timing influences functional durability. Regardless of whether NK cells were non-expanded or expanded for 10 or 17 days with or without feeders, all cultures ceased to proliferate within four weeks, indicating that earlier harvest does not extend the overall proliferative window – it simply accelerates cell division during the initial phases. This pattern suggests that cryopreservation may reset the proliferation “countdown,” leading to similar long-term behavior irrespective of expansion duration. Notably, NK cells from all time points remained responsive to re-stimulation eight weeks into long-term culture, indicating that NK cells retain the ability to divide upon encountering suitable target cells after extended resting periods. This is encouraging for NK cell-based immunotherapies and suggests that expanded NK cells may retain functional resilience *in vivo* [55].

Given current evidence that NK cell persistence is limited compared to T cells, repeated dosing is likely necessary regardless of expansion strategy [31,56,57]. This supports the use of later harvests, which maximize cell yield and reduce manufacturing cost per dose. As manufacturing platforms evolve, optimizing harvest timing –possibly even further than two weeks – will remain a key lever in balancing bioprocess efficiency, product longevity, and clinical dosing requirements.

### Cryopreservability of feeder cells and NK cells is essential for manufacturing

From a manufacturing perspective, the ability to cryopreserve feeder cells as standardized reagents and expanded NK cells as a therapeutic product without loss of function represents a significant advantage [31,58]. We observed that cryopreservation at high density and thawing without centrifugation improves NK cell yield after the thawing process. NK cells also retained a proliferation boost post-thaw and responded robustly to IL-2 stimulation, which is encouraging as IL-2 is known to enhance NK cell cytotoxicity, partly through upregulation of activating receptors such as NKp30, which plays a key role in tumor recognition [59]. Pre-activation of therapeutic NK cells before infusion could therefore be beneficial, paving the way for flexible, scalable, and potentially cost-effective NK cell therapy platforms. Looking ahead, GMP-compati le “thaw-and-activate” NK cell workflows and infusion bag formats that allow controlled addition of warm oxidized media and supplements may further improve therapeutic performance without compromising chain-of-custody in clinical settings.

### Future industry directions

Several practical and biological challenges remain when translating these laboratory findings into production for clinical use. While master cell banks of feeder cells can be generated in GMP-facilities, and γ-irradiated feeder cells can be characterized and cryopreserved, their use as GMP-grade reagents requires careful validation and regulatory oversight.

Next feeder-based expansion research should focus on setting up standardized methods evaluating and characterizing feeder cell reagents as well as final NK cell products. E.g. proliferative potential of γ-irradiated feeder cells, endotoxins, sterility (no bacteria, fungus, mycoplasma contamination), and feeder cell clearance validations (e.g. GFP+ feeder cells seen with flowcytometry and microscopy, or no dead feeder cell remnants like GFP’s DNA detected with qPCR) [60,61]. Furthermore, rapidly expanding NK cells can quickly outgrow their culture environment leading to arrested proliferation – underscoring the need for NK cell-optimized bioreactor systems. At scale, bioreactor choice should prioritize robust gas exchange and media supplementation to support rapid cell division and closed-system operation with automated monitoring.

Functionally, our study revealed that NK cells expanded under different conditions exhibit different cytotoxic traits. Future NK cell studies should explore how expansion conditions influence NK cell pharmacokinetics (e.g migration, persistence and clearance), target selection and responsiveness *in vivo*, as well as standardized NK cell potency assays for final product control. In conclusion, this study provides a comprehensive strategy for optimizing feeder-based expansion for scaled-up NK cell production. While further advances on creating more efficient genetically engineered NK cells will be essential in realizing NK cells full therapeutic potential – they will require robust feeder-based expansion technology to be utilized also economically [21].

## CONCLUDING REMARKS

Feeder-based expansion provides a practical and highly effective route to producing large and consistent NK cell batches required for off-the-shelf immunotherapies. This study identifies feeder cell designs, especially K562-mbIL21-CD48-41BBL, that stimulate NK cells to divide rapidly, with strong post-thaw proliferation, functional longevity and cytotoxicity – traits associated with therapeutic performance. These insights offer design principles for manufacturers and help bridge immunological discovery with industrial cell-therapy engineering. By outlining translational considerations such as optimal harvest timing, cryopreservation and thawing performance, phenotype-guided product matching and target-dependent cytotoxic profiles, our work supports the development of standardized, high-performance NK cell manufacturing systems.

Looking forward, feeder cell technologies can be positioned as core bioprocess reagents for NK cell manufacturing. It will however remain to be seen if NK cells from different sources (e.g. peripheral vs cord blood) would benefit from differently stimulating feeder cells – and should the second round of feeder cells be different than used in the first round. Further design in feeder cells could also be used to support different immunotherapy applications.

Currently there is limited information available on how feeder cells have been used to produce NK cells for clinical trials. This makes it difficult to transparently evaluate best methods, bioreactors and realistic yield expectations for NK cell manufacturing. GMP-grade feeder cell reagents could however enable reproducible manufacturing across sites with algorithm-driven automated process control. Near-future priorities include characterization of feeder cells, validated feeder cell removal, automated bioreactor operation, and harmonized release testing that links phenotype to mechanism of action. Similarly, more research and engineering efforts should be made towards improving the cryopreservation-thawing-activation phase, which will determine NK cells immediate survivability after infusion into patients. Together, these advances have the potential to reduce cost per dose, improve workflow efficiency, and collectively accelerate translation to consistent off-the-shelf clinical NK cell products supporting wide range of cancer treatments.

## METHODS

### Cell lines

The K562 chronic myelogenous leukemia cell line was purchased from ATCC (K-562; CCL-243) and used to generate the modified feeder cells. K562-mbIL15-41BBL feeder cells were a kind gift from Prof. Dario Campana*[16]*. K562 and all feeder cells were cultured in IMDM (Gibco, cat# 12440053) supplemented with 10% fetal bovine serum (Gibco, cat# 10082-147) and 1% penicillin-streptomycin (Gibco, cat# 15140-122).

K562-luc2 cells (ATCC, cat# CCL-243) were cultured under the same conditions as K562 cells, with the addition of 8 µg/mL blasticidin (Sigma-Aldrich, cat# SBR00022) when cultured for longer than one week.

NALM-6-Red-F-luc-GFP cells were previously created [5]. Briefly, NALM-6 cells (ATCC, cat# CRL-3273) were transduced with the IVISbrite Red F-luc-GFP lentiviral vector (RediFect, PerkinElmer) at an MOI of 10. After transduction and expansion, GFP-positive cells were sorted using a Sony SH800 cell sorter (Sony Biotechnology, San Jose, CA, USA). Cells were cultured in RPMI-1640 medium (Gibco, cat# 52400-025) supplemented with 10% FBS, 1% penicillin-streptomycin, and 1% L-glutamine (Gibco, cat# 25030-081).

SH-Sy5y-eGFPluc cells were also previously created *[62]*. Briefly, DNA encoding enhanced green fluorescent protein (eGFP) and firefly luciferase (GenBank ID: M15077.1), separated by a P2A ribosomal skip sequence, was synthesized and cloned into a pLV plasmid. Third-generation lentiviral vectors carrying the luciferase-eGFP gene were produced and used to transduce SH-Sy5y cells. Expression of eGFP and luciferase was confirmed by flow cytometry. Cells were cultured in EMEM (ATCC, cat# 30-2003) supplemented with 50% F12 Nutrient Mix (Thermo Fisher Scientific, cat# 11765-054), 10% FBS, 1% penicillin-streptomycin, and 1% L-glutamine.

### Gene design and viral vector production

New transgenes for feeder cells were designed *in silico* using protein sequences from the UniProt database *[63]*. DNA sequences were reverse-translated from protein sequences using the EMBOSS Backtranseq tool *[64]*. Transgene design was performed using Benchling (Benchling [Biology Software], 2021).

Transgenes were based on sequences from human CD8α (UniProt:P01732), IL-21 (Q9HBE4), IL-12A (P29459), IL-12B (P29460), IL-18 (Q14116), properdin (P27918), and 4-1BBL (P41273), fused together by linker sequences.

- mbIL21: Constructed using the CD8α signal peptide (aa 1–21), IL-21 domain (aa 25–162), a (GGGGS)_2_ linker, and CD8α hinge (aa 136 –182), transmembrane (aa 183–203), and cytoplasmic regions (aa 204– 213).
- mbIL12-18-21: Included CD8α signal peptide, IL-12B/IL-12p40 (aa 23–328), IL-12A/IL-21p35 (aa 23– 219), IL-18 (aa 37–193), and IL-21 (aa 25–162) domains, separated by flexible linkers, anchored with CD8α hinge, transmembrane and cytoplasmic regions.
- Mbproperdin: Included CD8α signal peptide, a myc tag (EQKLISEEDLLRKRRT), a flexible G4S based linker, and the properdin domain (aa 28–469), fused to CD8α hinge, transmembrane and cytoplasmic regions.
- mbCD48: Included CD8α signal peptide and CD48 domain (aa 27– 220), linked to CD8α hinge,transmembrane and cytoplasmic regions.
- 4-1BBL: Contained the full-length 4-1BBL sequence (aa 1–254).cDNAs were synthesized and cloned into lentiviral transfer plasmids by GENEWIZ (Azenta Life Sciences). Third-generation vesicular stomatitis virus glycoprotein G (VSV-G) pseudotyped lentiviral vectors, were produced at the National Virus Vector Laboratory, A.I. Virtanen Institute for Molecular Sciences, University of Eastern Finland *[65,66]*.

### Transduction and irradiation of feeder cells

K562 feeder cells expressing mbIL21, mbIL12-18-21, mbProperdin, mbCD48, 4-1BBL, and GFP were generated by transducin K5 cells with lentiviral vectors at an I of 5. Cells were transduced overni ht at a density of 1×10^6^ cells/ml, then washed and cultured at 0,1×10^6^ cells/ml.

Transduction efficiency was assessed by flow cytometry using non-transduced cells as negative controls. Antibodies are listed in Supplementary Table 2. K562, K562-mbIL21, K562-mbIL21-CD48, and K562-mbIL21-CD48-41BBL feeder cells (all GFP-positive) were sorted based on IL-21, CD48, 4-1BBL, and GFP expression the Biomedicum FACS Core Facility, University of Helsinki.

Feeder cells were gamma-irradiated with 100 Gy using a gamma irradiator (OB29/4, STS, Braunschweig, Germany) with a Cs-137 source. Irradiated cells were cryopreserved in culture medium containing 5% DMSO (Wak Chemie, cat# 281235).

### NK cell expansion

On day 0, peripheral blood was donated by healthy volunteers at the Finnish Red Cross Blood Service. On day 1, primary human peripheral blood mononuclear cells (PBMCs) were isolated from buffy coats obtained from these donations at the Finnish Red Cross Blood Service under institutional permit FRCBS 178/6/2023. PBMCs were separated by density gradient centrifugation using Ficoll-Paque (GE Healthcare, cat# 17-1440-02). NK cells were isolated using the NK Cell Isolation Kit (Miltenyi Biotec, cat# 130-092-657) accordin to the manufacturer’s instructions and cryopreserved in MACS Freezing solution (Miltenyi Biotec, cat# 130-129-552). Purity of PBMCs and NK cells was assessed by flow cytometry with FACSymphony A1 Cell Analyzer (BD Biosciences). Antibodies and isotype controls are listed in Supplementary Table 2.

Thawed NK cells were cultured in NK MACS Expansion Medium consisting of NK MACS basal medium (Miltenyi Biotech, NK MACS medium, cat# 130-112-968) supplemented with 1 % NK MACS supplement (Miltenyi Biotech, NK MACS supplement, cat# 130-113-102), 5 % human AB serum (MP Biomedicals, Human AB Serum, cat# 2930949 or Sigma, Human AB Serum, cat# H5667), 500 IU/mL IL-2 (Miltenyi Biotech, Human IL-2 IS premium grade cat# 130-097-746), 140 IU/ml IL-15 (Miltenyi Biotech, Human IL-15 premium grade, cat# 130-095-765) 50 U/ml penicillin and 50 g/ml streptomycin (Gibco, Penicillin-Streptomycin, cat#15140-122).

On day 3, 1×10^5^ NK cells were stimulated with 5×10^5^ irradiated feeder cells. Cultures were monitored, counted and subcultured as needed. On day 10, 1×10^6^ NK cells were re-stimulated with 5×10^6^ feeder cells. On day 17, NK cells were collected and cryopreserved for later assays.

Cumulative population doubling (cPD) of NK cells during culture was calculated using the following formula:

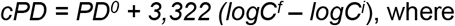

where

PD^0^ is previous population doubling

logC^f^ is final number of cells in culture

logC^i^ is starting number of cells in culture

### DNA isolation and telomere length assay

DNA was extracted from isolated (day 1) and expanded (day 17) NK cells immediately after thawing using the DNeasy Blood & Tissue Kit (QIAGEN, cat# 69505) and RNase (QIAGEN, cat# 19101), following the manufacturer’s instructions. DNA concentration was measured using a NanoDrop 2000 spectrophotometer (Thermo Scientific). DNA samples were stored at − 80 °C and analyzed for telomere length.

Measurement of relative telomere length (rTL) from NK cells was performed using quantitative PCR (qPCR). DNA from day 1 and 17 samples were used for the telomere and *HBB* (reference single copy gene) qPCRs. Altogether 25 ng of DNA from each sample were run in triplicates on 384-well plates with CFX 384 PCR system (BioRad, Hercules, CA).

The qPCR primer sequences were

Telomere_F: CGGTTTGTTTGGGTTTGGGTTTGGGTTTGGGTTTGGGTT;

Telomere_R: GGCTTGCCTTACCCTTACCCTTACCCTTACCCTTACCCT;

HBB_F: GCTTCTGACACAACTGTGTTCACTAGC; and

HBB_R: CACCAACTTCATCCACGTTCACC,

and the qPCR conditions for both telomere and *HBB* followed a previous protocol*[67]*. Previously assayed DNA sample extracted from whole blood was included as a plate and assay control. rTL was calculated by subtracting telomere qPCR cycle threshold (C_t_) from the *HBB* qPCR C_t_ (ΔC_t_). Telomere change between the time points (at day17 vs. day 1) was calculated as percentage change: (2^ (ΔC_t_ at time point 2 – ΔC_t_ at time point1)-1*100).

### Long term culture

Thawed NK cells were cultured in modified NK MACS Expansion Medium, where IL-2 concentration was reduced from 500 IU/ml to 100 IU/ml and penicillin-streptomycin replaced with 1X MycoZap-Plus-PR (Lonza, cat# LZ-VZA-2021). Cell count and complete medium replacement with centrifugation of cells at 200 x g for 7 min was performed once weekly. Every week, a new culture was started with a maximum of 4×10^6^ cells/sample. The long-term culture was continued until the cell number was reduced below 0,2×10^6^ cells/sample, after which cultures were considered dead and the culture ended.

### Cytotoxicity assay

Unless used immediately, NK cells were cultured in NK MACS Expansion Medium. Prior to the assay, NK cells were transferred to cytokine-free NK MACS Expansion Medium. Target cells (5 000 cells/well) and NK cells (at various effector-to-target [E:T] ratios) were co-cultured in 384-well plates in total volume of 30 µl/well, using a 1:1 mixture of cytokine-free NK MACS Expansion Medium and the respective target cell medium. For the IL-2 responsiveness assay, 500 IU/ml of IL-2 was added to the co-culture medium.

Co-cultures were incubated for 24 hours at 37°C with 5% CO_2_, after which 25 µl/well luciferase assay reagent (Promega, ONE-Glo Luciferase assay system, cat# E6110) was added and the remaining live target cells were quantified with a CLARIOstar microplate reader (BMG Labtech).

### Phenotyping

Phenotyping was performed on isolated and expanded NK cells five days post-thawing. During the recovery period, NK cells were cultured in NK MACS Expansion Medium. Cells were then labeled and analyzed by flow cytometry. Antibodies and isotype controls are listed in Supplementary Table 2.

### Degranulation assay

Degranulation and cytokine secretion assays were performed similarly to the cytotoxicity assay. NK cells were cultured in NK MACS Expansion Medium for seven days post-thawing, then washed and transferred to cytokine-free NK MACS Expansion Medium.

A total of 150 000 NK cells were cultured with or without 300 000 K562 target cells in 200 µL of medium. Anti-CD107a antibody or isotype control was added at the start of incubation. After 1 hour, a protein transport inhibitor (Protein Transport Inhibitor Cocktail, Invitrogen, cat# 00-4980) was added. Cells were co-cultured for a total of 4 hours, after which samples were collected for labeling and flow cytometry analysis. Antibodies and isotype controls are listed in Supplementary Table 2.

For secreted cytokine analysis, culture supernatants were collected without transport inhibitor treatment and stored at − 80 °C for multiple analysis.

### Flow cytometry

Flow cytometry was used to analyze expression of the transgenes on feeder cells, the purity of isolated NK cells and PBMCs, and phenotype and activation markers of NK cells. Cells were washed with flow cytometry buffer (0.5% human serum albumin [Albunorm] in PBS), blocked with Fc-blocking solution (BioLegend, Human TruStain FcX, cat# 422302), and surface-stained with antibodies and isotype controls (Supplementary Table 2). When required, cells were fixed with 4% paraformaldehyde (BioLegend, cat# 420801). Samples were analyzed using a Beckman Coulter DxFLEX or BD FACSymphony A1 Cell Analyzer. Data were processed with FlowJo Software version 10.8.1 (FlowJo LLC, Ashland, OR).

### Multiplex analysis of co-culture supernatants

Human cytokines IL-2, IL-4, IL-13, IFN-γ, N -α, ran yme and perforin were analy ed in a multiple manner y using MILLIPLEX MAP Human CD8+ T Cell Magnetic Bead Panel assay kit (Merck, Cat# HCD8MAG-15K). Cytokines were measured from undiluted cell supernatant samples. Single samples were analyzed using Bio-Plex 200 detector and the data was analyzed with Bio-Plex Manager 6.1 software using five-parameter logistic regression for standard curve fitting. The results are expressed as pg/ml based on the standard curve calculations. The lower limits of quantitation (LLOQ) were following: IL-2: 1.8 pg/ml, IL-4: 2.3 pg/ml, IL-13: 1.7 pg/ml, TNF-α 0.5p /ml, I N-γ 1. pg/ml, granzyme B: 1,2 pg/ml and perforin: 11.9 pg/ml.

### TGF-β1 ELISA

Cell culture supernatants were collected after 72 hours of NK cell expansion and stored at −80 °C. Prior to analysis, samples were thawed on ice and analyzed using the Human TGF-β1 DuoSet ELISA (Bio-Techne, cat# DY240-05) and DuoSet Ancillary Reagent Kit 1 (Bio-Techne, cat# DY007B), following the manufacturer’s instructions. Absorbance was measured using a CLARIOstar plate reader.

### CAR NK cell transduction and recovery from cryopreservation

Anti-CD19 FiCAR-NK1 (shortly CAR) construct design and transduction has been previously described in detail [5]. Briefly, Day 1 NK cells were cultured in NK MACS Expansion Medium. Three days later NK cells were transferred to NK MACS Expansion Medium supplemented with 5 µM BX-795 (MedChemExpress, cat# HY-10514), incubated for 30 min and plated on RetroNectin-coated wells: the wells had been coated earlier for 2 h with 25 µg/ml RetroNectin (Takara, cat# T100B) in PBS (Gibco), blocked 30 min with 2% BSA (Gibco, cat# 15260-037) and washed with PBS. On the plate, NK cells were transduced over night with VSV-G pseudotyped lentiviral vector (MOI 10) to express anti-CD19 FiCAR-NK1. Following day, CAR NK cells were washed and expanded for two weeks, with weekly K562-mbIL21-CD48-41BBL feeder cells stimulations at 1:2 NK:Feeder cell ratio.

After CAR NK cell expansion, cells were washed and resuspended at 10×10^6^ or 50×10^6^ cells/ml in five different cryopreservation media: 1) Albunorm 200g/l (Octapharma, cat# VNR054343) with 5% DMSO (Wak Chemie, cat# 281235), 2) Albunorm with 10% DMSO, 3) human AB serum (Sigma, cat# H5667) with 10% DMSO, 4) MACS Freezing solution (Miltenyi Biotech, cat# 130-129-552) or 5) complete NK MACS Expansion Medium with 50% Human AB Serum and 10% DMSO.

The frozen CAR NK cells were thawed either by direct addition to warm NK MACS Expansion Medium at 2×10^6^ cells/ml or by washing and centrifugation (300×g, 5 min, room temperature) before resuspension in warm medium. The Cells were cultured for 3 days post-thaw, and viability and cell counts were assessed immediately after thawing and after the 3-day culture period.

### Statistics and figures

Flow cytometry data were analyzed using FlowJo Software version 10.8.1. Gating strategies were standardized across samples, and results shown as quantitative summaries. Statistical analyses were performed using Microsoft Excel for Microsoft 365. Statistical significances were analyzed using a two-tailed paired, equal or unequal-variance t-tests. A p-value < 0.05 was considered statistically significant. Data are presented as mean ± standard deviation (SD) unless otherwise noted. Figures and data visualizations were generated using Microsoft Excel, and scientific illustrations were created using BioRender.

## RESOURCE AVAILABILITY

### Lead contact

Helka Göös, iCell Group, Research and Development, Finnish Red Cross Blood Service, Haartmaninkatu 8, FIN-00290 Helsinki, Finland, Finland. Tel. +358444344395

### Materials availability

Materials listed in the Key Resources Table were obtained from external providers as noted. Any new materials generated in this study are available from the lead contact upon reasonable request.

3Data and code availability

This paper dos not report original code. For original data, please contact (H. Göös).

## ACKNOWLEDGEMENTS

Acknowledgements

This research was conducted at the Finnish Red Cross Blood Service and the University of Eastern Finland. We gratefully acknowledge funding from the Research Council of Finland GeneCellNano Flagship Program and Orion Corporation. We thank the FACS Core Facility at Biomedicum, University of Helsinki, for their expert services. We are especially grateful to Anne Martikainen from the National Virus Vector Laboratory, A.I. Virtanen Institute, University of Eastern Finland, for producing and titrating the lentiviral vectors used in this study. We also thank Laurens Veltman, MSc, for research assistance and Anu Autio, PhD, for valuable feedback and comments. Finally, we thank Dario Campana, MD, PhD, for generously providing us with feeder cells.

## Authorcontributions

M.S., F.J., K.V., H.G. and M.K. conceived and designed the study. F.J., H.P., M.K., K.V. and S.Y-H. supervised the study and coordinated funding. M.S. performed *in silico* design. M.S., F.J. and E.K. designed flow cytometry panels. M.S. and F.J. designed the experiments. M.S., F.J. and L.A. performed the experiments. M.S. and F.J. analyzed the data. H.G. performed and analyzed cryopreservation recovery experiment. J.K. and J.E. contributed to the experiments. A.S. and M.S. performed and analyzed telomere length measurement. D.S. coordinated and designed lentiviral vector production. P.V-K. performed multiplexing experiments. M.S. wrote the manuscript. All authors reviewed the manuscript and approved the final version.

## GLOSSARY

ADCC (antibody-dependent cellular cytotoxicity): A killing mechanism where NK-cell receptor CD16 binds antibody-coated targets
Activating ligand: A molecule on target cells that engages activating receptors
Adaptive immunity: Antigen-specific immunity involving T and B cells
Allogeneic: Originating from a genetically different donor of the same species
CAR (chimeric antigen receptor): An engineered receptor giving immune cells new target specificity
CD3 (cluster of differentiation 3): A T-cell marker; its absence (CD3–) distinguishes NK cells
CD16 (cluster of differentiation 16 / FcγRIII): An Fc receptor mediating ADCC and identifying cytotoxic NK subsets
CD56 (cluster of differentiation 56 / NCAM): Canonical NK-cell marker used to identify NK cells and define subsets
CD57 (cluster of differentiation 57): A terminal maturation marker of NK cells
CD69 (cluster of differentiation 69): An early activation marker upregulated after stimulation
CD107a (cluster of differentiation 107a / LAMP-1): A degranulation marker used to measure NK-cell cytotoxicity
Cell sorting: A technique used to isolate specific cells using fluorescence or physical traits
Chimeric transgene construct: An engineered sequence combining multiple genetic elements
Co-stimulatory ligand: Provides additional activation signals to immune cells
Cord blood: Umbilical-cord-derived blood rich in stem and immune cells
CRS (cytokine release syndrome): A systemic inflammatory reaction caused by excessive cytokine secretion
Cryopreservation: Freezing cells for long-term storage
Cumulative proliferation curve: A graph showing total cell expansion over time
Cytokine: A protein that mediates immune communication
Cytokine secretion: Release of cytokines following activation
Cytotoxic granule: Vesicles containing perforin and granzymes
Cytotoxicity: The ability of immune cells to kill target cells
Degranulation: Release of cytotoxic granules during target-cell killing
Effector cell: An immune cell executing functions such as killing or cytokine production
Epitope masking: Concealment of antigenic sites reducing recognition
Ex vivo: Performed on cells outside the body
Exhaustion: Dysfunction caused by chronic stimulation
FasL (Fas ligand): A death-inducing ligand triggering apoptosis through the Fas receptor
Feeder cell: Cells providing stimulatory signals for immune-cell expansion
Flow cytometry: Fluorescence-based technique for characterizing cells
Fratricide: Immune cells killing each other due to shared antigens
GFP (green fluorescent protein): Afluorescent reporter used to track gene expression or label cells
Granzyme B: A cytotoxic enzyme inducing apoptosis in target cells
GvHD (graft-versus-host disease): A reaction where donor immune cells attack host tissues
IFN-γ (interferon gamma): A cytokine that boosts inflammatory and immune activation
Immunosuppressive: Capable of dampening immune responses
Immunotherapy: Treatments that harness or modify the immune system
In silico: Computer-based analysis
In vitro: Performed outside the organism in controlled environments
In vivo: Performed within a living organism
Interleukin: A cytokine family regulating immune function
K562 (human chronic myelogenous leukemia cell line): A standard target line in cytotoxicity assays and classic feeder line for NK-cell expansion
LAG-3 (lymphocyte activation gene-3): An inhibitory receptor involved in immune-cell exhaustion
mb (membrane-bound): Describes molecules anchored to the cell membrane
Memory-like: NK cells with durable, enhanced functional capacity
MFI (median fluorescence intensity): A flow-cytometry measure of marker expression
NALM-6 (B-cell acute lymphoblastic leukemia cell line): A standard target line in cytotoxicity assays
NK (natural killer): An innate lymphocyte responsible for cytotoxic and cytokine-mediated immunity
NKG2A (NK group 2A): An inhibitory receptor recognizing HLA-E
NKG2D (NK group 2D): An activating receptor recognizing stress-induced ligands
PBMCs (peripheral blood mononuclear cells): Blood-derived lymphocytes and monocytes
Perforin: Pore-forming protein enabling granzyme delivery into target cells
Phenotype: Observable traits or marker expression profiles
Primary NK cell: A donor-derived NK cell directly isolated from blood
Proliferation: Cell division and expansion
Senescence: A growth-arrested state associated with reduced cellular function
SH-SY5Y (human neuroblastoma cell line): A tumor cell line used in cytotoxicity assays
Surface expression: Presence of specific proteins on the cell membrane
TGF-β1 (transforming growth factor beta-1): An immunosuppressive cytokine inhibiting NK-cell activation
TIGIT (T-cell immunoreceptor with Ig and ITIM domains): An inhibitory receptor competing with DNAM-1 for shared ligands
Telomere: Protective DNA structures at chromosome ends that shorten with division
TNF-α (tumor necrosis factor alpha): A pro-inflammatory cytokine produced during immune activation
Transduction: Viral delivery of genetic material into cells
TRAIL (TNF-related apoptosis-inducing ligand): A ligand triggering apoptosis through death receptors

