## Supplemental Information for "Feeder cell – the key component in producing scalable and fit NK cells for therapeutic use"

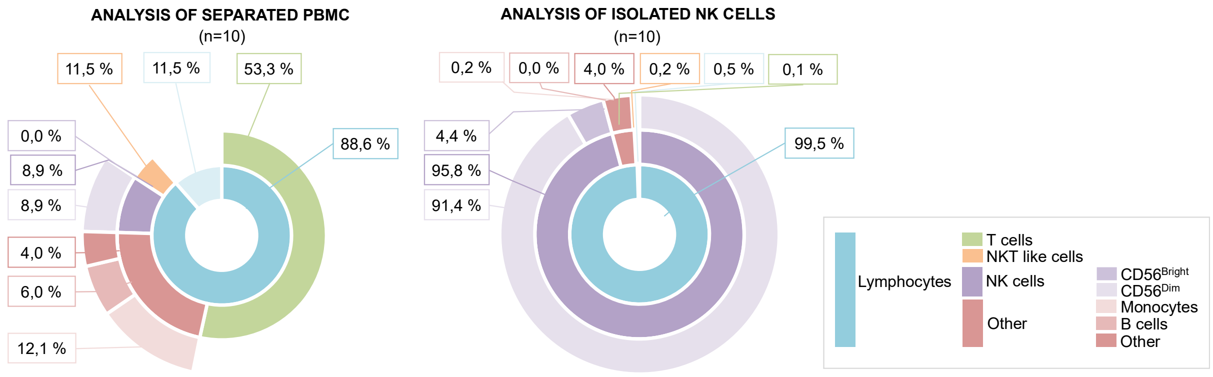


**Supplementary Figure 1.** **Purity analysis of PBMCs and isolated NK cells.** Sunburst chart illustrates the composition of immune cell populations in buffy coat-derived peripheral blood mononuclear cells (PBMCs) and NK cell-enriched fractions following isolation. The charts display the relative proportions of lymphocytes, T cells, NKT-like cells, NK cells, B cells, and monocytes within each sample type, based on flow cytometry analysis. Data are presented as mean from n = 10 individual donors.

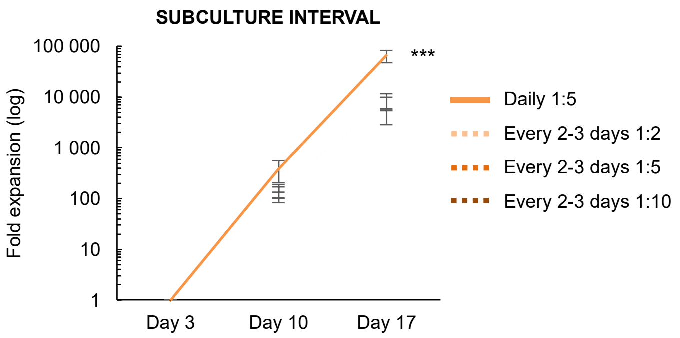


**Supplementary Figure 2.** **Subculture interval affects NK cell expansion.** Impact of subculture intervals and effector-to-target (E:T) ratios on NK cell fold expansion using highly stimulatory K562-mbIL21-CD48-41BBL feeder cells. Data represents cumulative expansion over the culture period under different conditions. Data are presented as mean ± SD from 3 individual donors. Statistical analysis comparison to Every 2-3 days 1:2 was determined using a two-tailed unequal-variance t-test. $p < 0.05$ (*), $p < 0.01$ (**), $p < 0.001$ (***).

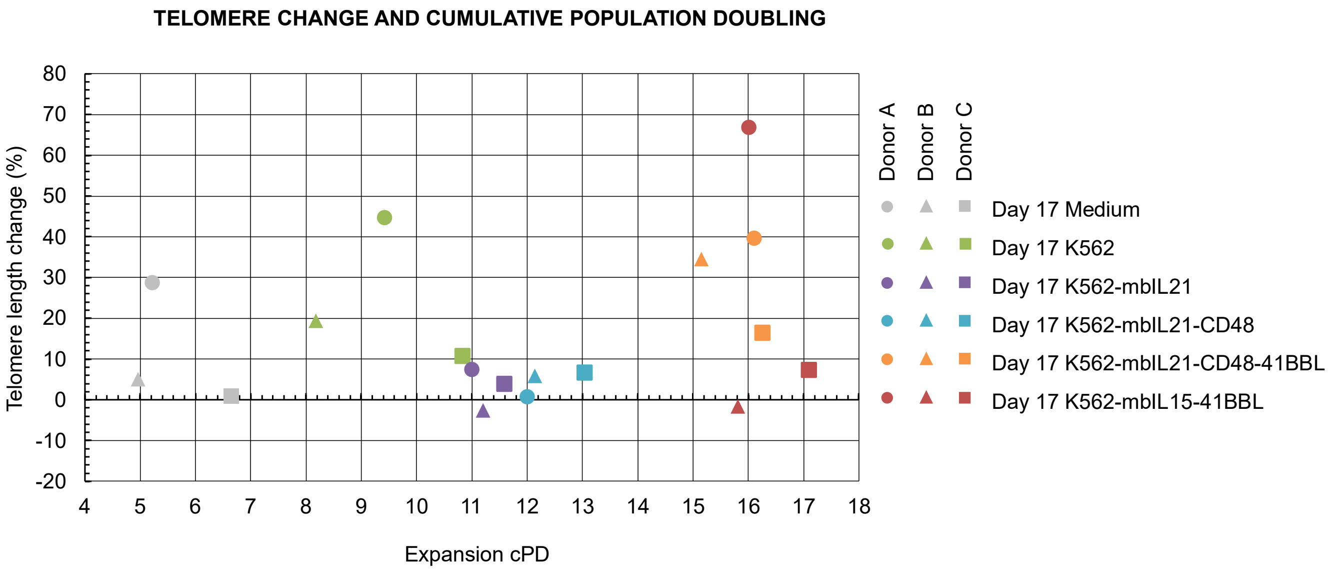


**Supplementary Figure 3. Expansion efficiency and purity of NK cells expanded with or without feeder cells.** Relationship between NK cell expansion and telomere length dynamics during production with selected feeder cells. DNA was isolated from NK cells at the beginning of the culture (Day 1) and after 17 days of expansion. Telomere length was measured using a PCR-based method, and the relative change over time was calculated for each donor. These changes are shown in relation to the expansions cumulative population doubling (cPD) achieved during the same period. Each data point represents an individual donor: donor A (circles), B (triangles) and C (squares). Data is presented from n = 3 individual donors.

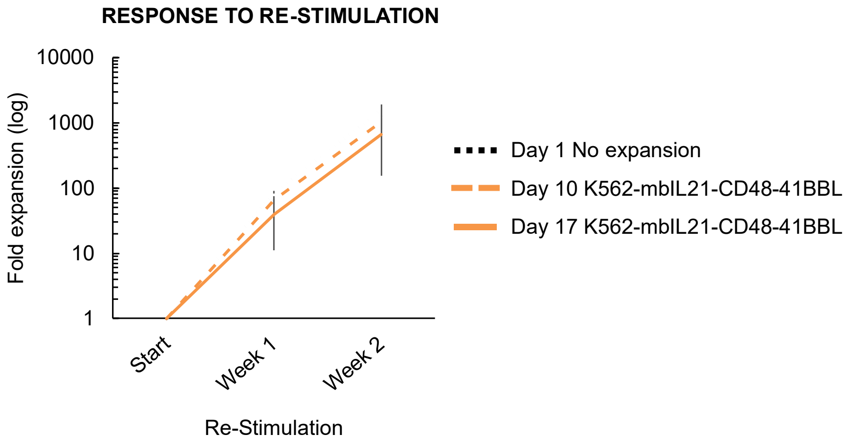


**Supplementary Figure 4. Capacity of expanded NK cells for re-stimulation after 8 weeks of long-term *in vitro* culture.** After eight weeks, selected NK cell populations were re-stimulated with K562-mbIL21-CD48-41BBL feeder cells for 2 weeks to assess retained proliferative capacity. Data are presented as mean ± SD from 3 individual donors. Statistical analysis comparisons were made against the Day 1, non-expanded NK cell condition unless otherwise indicated using two-tailed paired t-test. p < 0.05 (*), p < 0.01 (**), p < 0.001 (***).

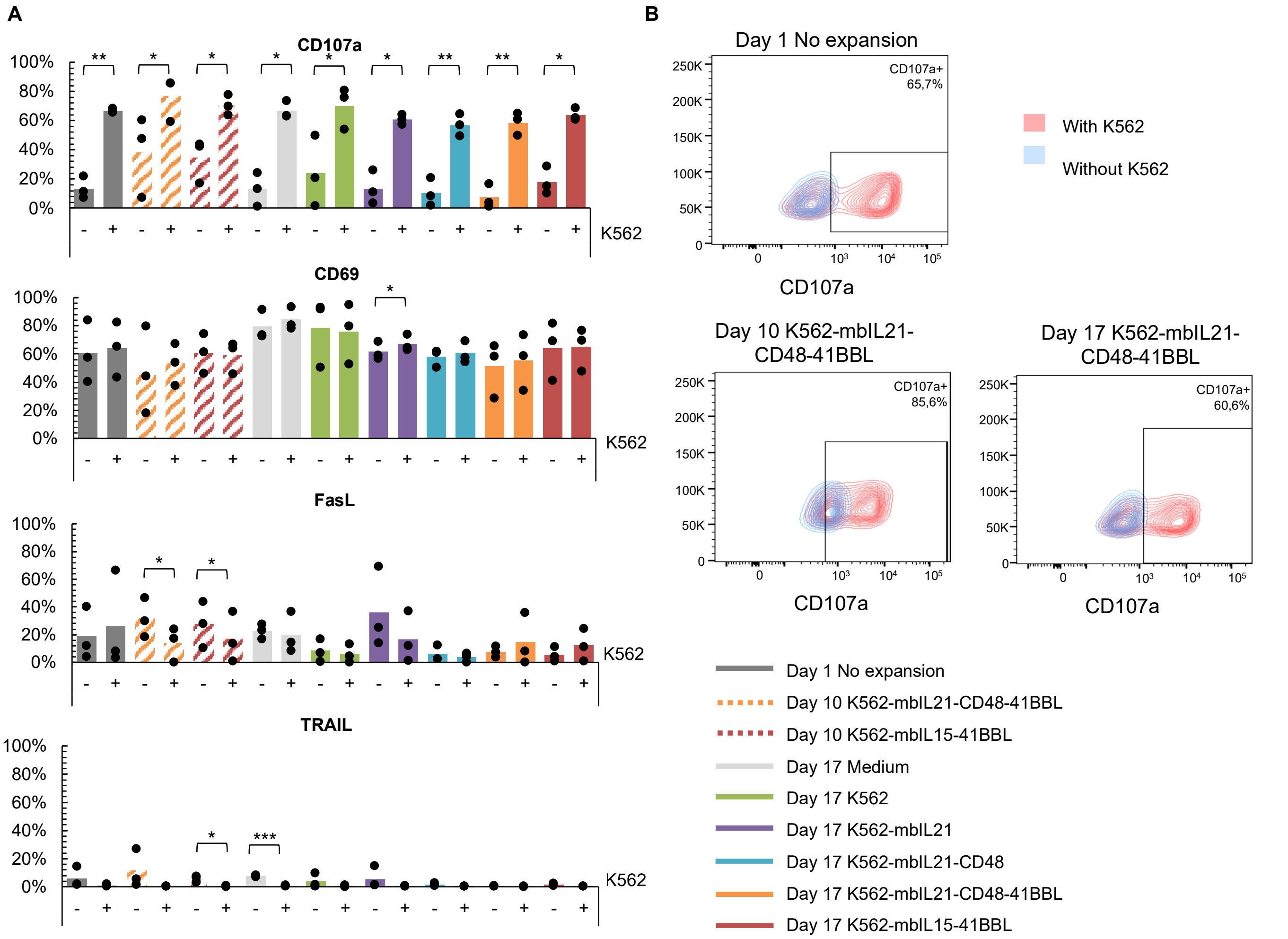


**Supplementary Figure 5. Activation marker expression of expanded NK cells in response to K562 target cells.** (A) NK cells were co-cultured for 4 hours with (+) or without (–) K562 target cells and analyzed by flow cytometry. Surface expression of the activation marker CD69, degranulation marker CD107a, and apoptosis-inducing ligands FasL and TRAIL were measured on NK cells. (B) Representative flowcytometry plots showing degranulation of NK cells at different stages of expansion. Data are presented as mean ± SD from (A) n = 3 individual donors. Data (A) points represent individual donors: donor A (circles), B (triangles) and C (squares). Statistical analysis comparisons were made against the Day 1, non-expanded NK cell condition unless otherwise indicated using two-tailed paired t-test. $p < 0.05$ (*), $p < 0.01$ (**), $p < 0.001$ (***).

**Supplementary Table 1. Summary of NK cell expansion and population doubling under different expansion conditions.** NK cells were cultured with or without various feeder cell types at 1:5 ratio and monitored over a 14-day period from Day 3 to 17. Fold expansion and population doubling times were quantified at Days 10 and 17 of expansion, with data presented as mean of n=3-6 individual donors ± SD. Statistical significance comparing different conditions to Medium expansion was determined using two-tailed paired t-test. $p < 0.05$ (*), $p < 0.01$ (**), $p < 0.001$ (***).

| NK cell expansion condition | Fold expansion | | Cumulative population doublings | | |
|---|---|---|---|---|---|
| | Day 10 | Day 17 | Day 10 | Day 17 | |
| Medium (n=6) | 11 ± 9 | 160 ± 157 | 3,1 ± 1,1 | 6,7 ± 1,5 | - |
| K562 (n=6) | 46 ± 27 | 1 162 ± 836 | 5,3 ± 1,0 | 9,7 ± 1,4 | * |
| K562-mbProperdin (n=3) | 30 ± 19 | 1 208 ± 1 102 | 4,7 ± 1,2 | 9,4 ± 2,4 | ns |
| K562-mbCD48 (n=3) | 29 ± 15 | 1 299 ± 970 | 4,7 ± 1,0 | 9,8 ± 1,8 | ns |
| K562-mbIL12-18-21(n=3) | 45 ± 32 | 1 781 ± 1 774 | 5,0 ± 1,6 | 9,8 ± 2,5 | ns |
| K562-mbIL21 (n=6) | 77 ± 44 | 3 924 ± 3 135 | 6,0 ± 1,0 | 11,5 ± 1,3 | * |
| K562-mbIL21-Properdin (n=3) | 58 ± 38 | 4 758 ± 3 822 | 5,5 ± 1,4 | 11,4 ± 2,3 | ns |
| K562-mbIL21-CD48 (n=6) | 88 ±51 | 5 820 ± 3 165 | 6,2 ± 1,0 | 12,2 ± 1,2 | ** |
| K562-41BBL (n=3) | 112 ± 31 | 22 731 ± 9 297 | 6,8 ± 0,4 | 14,4 ± 0,7 | ns |
| K562-mbIL21-41BBL (n=3) | 217 ± 55 | 41 680 ± 16 165 | 7,7 ± 0,4 | 15,3 ± 0,7 | * |
| K562-mbIL21-CD48-41BBL (n=6) | 387 ± 181 | 65 939 ± 17 854 | 8,5 ± 0,6 | 16,0 ± 0,5 | *** |
| K562-mbIL15-41BBL (n=6) | 337 ± 108 | 82 267 ± 38 084 | 8,3 ± 0,4 | 16,2 ± 0,7 | ** |

**Supplementary Table 2.** **List of antibodies and isotype controls used in flow cytometry analysis.**

| REAGENT | MANUFACTURER | CLONE | CATALOG NUMBER |
|---|---|---|---|
| 7-AAD | Miltenyi Biotec | - | 170-081-088 |
| PE-Dazzle 594 anti-human CD107a (LAMP-1) antibody | BioLegend | H4A3 | 328646 |
| PerCP/Cy5.5 anti-human CD159a (NKG2A) antibody | BioLegend | S19004C | 375126 |
| Brilliant Violet anti-human CD16 antibody | BioLegend | 3G8 | 302038 |
| APC anti-human CD178 (Fas-L) antibody | BioLegend | NOK-1 | 306421 |
| PE/Cy7 anti-human CD253 (Trail) antibody | BioLegend | RIK-2 | 308216 |
| PE/Dazzle 594 anti-human CD314 (NKG2D) antibody | BioLegend | 1D11 | 320828 |
| PE anti-human CD57 recombinant antibody | BioLegend | QA17A04 | 393308 |
| PE anti-human IL-12/IL-23 p40 | InVitrogen | eBioHP40 | 12-7235-42 |
| APC/Cy7 anti-human TIGIT (VSTM3) antibody | BioLegend | A15153G | 372734 |
| APC anti-human CD3 antibody | BioLegend | 6K7 | 344812 |
| APC Mouse IgG1, κ Isotype Ctrl (FC) Antibody | BioLegend | MOPC-21 | 400122 |
| APC/Cyanine7 Mouse IgG2a, κ Isotype Ctrl Antibody | BioLegend | MOPC-21 | 400230 |
| Brilliant Violet 421 anti-human CD137L (4-1BBL) antibody | BioLegend | 5F4 | 311508 |
| Brilliant Violet 421 Mouse IgG1, κ Isotype Ctrl Antibody | BioLegend | MOPC-21 | 400158 |
| Brilliant Violet 510 anti-human CD56 (NCAM) ntibody | BioLegend | HCD56 | 318340 |
| Brilliant Violet 510 Mouse IgG1, κ Isotype Ctrl Antibody | BioLegend | MOPC-21 | 400172 |
| Brilliant Violet 605 anti-human CD56 (NCAM) | BioLegend | HCD56 | 318334 |
| Brilliant Violet 605 Mouse IgG1, κ Isotype Ctrl Antibody | BioLegend | MOPC-21 | 400162 |
| Brilliant Violet 785 anti-human CD69 antibody | BioLegend | FN50 | 310932 |
| Brilliant Violet 785 anti-human CD223 (LAG-3) | BioLegend | 11C3C65 | 369322 |
| Brilliant Violet 785 Mouse IgG1, κ Isotype Ctrl Antibody | BioLegend | MOPC-21 | 400170 |
| CD14 antibody, anti-human, APC, REAfinity | Miltenyi Biotec | REA599 | 130-110-578 |
| CD19 antibody, anti-human, PE-Vio770, REAfinity | Miltenyi Biotec | REA675 | 130-114-173 |
| CD3 antibody, anti-human, FITC, REAfinity | Miltenyi Biotec | REA613 | 130-113-700 |
| CD45 antibody, anti-human VioBlue, REAfinity | Miltenyi Biotec | REA747 | 130-110-775 |
| CD56 antibody, anti-human, PE, REAfinity | Miltenyi Biotec | REA196 | 130-113-876 |
| Complement factor P (Properdin) monoclonal antibody | InVitrogen | KT20 | MA172515 |
| Alexa Fluor 488 Human IL-18/IL-1F4 | BioTechne | 925008 | C2548G |
| PE IL-12 p35 monoclonal antibody | InVitrogen | 27537 | MA523559 |
| PE anti-human IL-21 antibody | BioLegend | 3AE-N2 | 513004 |
| PE Mouse IgG1, κ Isotype Ctrl (FC) Antibody | BioLegend | MOPC-21 | 400114 |
| PE/Cyanine7 anti-human CD48 | BioLegend | BJ400 | 336718 |
| PE/Cyanine7 Mouse IgG1, κ Isotype Ctrl Antibody | BioLegend | MOPC-21 | 400126 |
| PE/Dazzle™ 594 Mouse IgG1, κ Isotype Ctrl Antibody | BioLegend | MOPC-21 | 400176 |
| PerCP/Cyanine5.5 Mouse IgG1, κ Isotype Ctrl Antibody | BioLegend | MOPC-21 | 400150 |
| PE Rat anti-mouse IgG1 secondary antibody | InVitrogen | m1-14D12 | 12-4015-82 |
| REA Control Antibody (S), human IgG1, APC, REAfinity | Miltenyi Biotec | REA293 | 130-113-434 |
| REA Control Antibody (S), human IgG1, FITC, REAfinity | Miltenyi Biotec | REA293 | 130-113-437 |
| REA Control Antibody (S), human IgG1, PE, REAfinity | Miltenyi Biotec | REA293 | 130-113-438 |
| REA Control Antibody (S), human IgG1, PE-Vio 770, REAfinity | Miltenyi Biotec | REA293 | 130-113-440 |
| REA Control Antibody (S), human IgG1, VioBlue, REAfinity | Miltenyi Biotec | REA293 | 130-113-442 |
| Zombie Green | BioLegend | - | 423111 |
